# Stroke-related alterations in inter-areal communication

**DOI:** 10.1101/2021.01.04.425190

**Authors:** Michele Allegra, Chiara Favaretto, Nicholas Metcalf, Maurizio Corbetta, Andrea Brovelli

## Abstract

Beyond causing local ischemia and cell damage at the site of injury, stroke strongly affects long-range anatomical connections, perturbing the functional organization of brain networks. Several studies reported functional connectivity abnormalities parallelling both behavioral deficits and functional recovery across different cognitive domains. FC alterations suggest that long-range communication in the brain is altered after stroke. However, standard FC analyses cannot reveal the directionality and time scale of inter-areal information transfer. We used resting-state fMRI and covariance-based Granger causality analysis to quantify network-level information transfer and its alteration in stroke. Two main large-scale anomalies were observed in stroke patients. First, inter-hemispheric information transfer was significantly decreased with respect to healthy controls. Second, stroke caused inter-hemispheric asymmetries, as information transfer within the affected hemisphere and from the affected to the intact hemisphere was significantly reduced. Both anomalies were more prominent in resting-state networks related to attention and language, and they correlated with impaired performance in several behavioral domains. Overall, our findings support the hypothesis that stroke provokes asymmetries between the affected and spared hemisphere, with different functional consequences depending on which hemisphere is lesioned.

## INTRODUCTION

Spontaneous brain activity is intrinsically organized into large-scale networks of correlated activity (Bullmore and Sporns, 2009; Damoiseaux et al., 2006; Fox et al., 2005), also known as resting-state networks (RSNs). The functional organization of RSNs is altered in stroke (Corbetta et al., 2018, 2015). In fact, local ischemia, which damages cells and neural connections at the site of injury, primarily affects subcortical regions and white matter, thus altering long-range functional connectivity (FC) between cortical areas. Two types of large-scale FC alterations affect RSNs (Joshua Sarfaty Siegel et al., 2016): i) a decrease of within-network interhemispheric FC (Carter et al., 2010; Golestani et al., 2013; He et al., 2007; New et al., 2015; Park Chang-hyun et al., 2011; Ramsey et al., 2016; Joshua Sarfaty Siegel et al., 2016; Tang et al., 2016); ii) an increase of between-network intra-hemispheric FC (Baldassarre et al., 2014; Eldaief et al., 2017; Ramsey et al., 2016; Joshua Sarfaty Siegel et al., 2016). As a consequence, within-RSN connections are weakened, while between-RSN connections are strengthened, which translates into an overall decrease of network modularity (Gratton et al., 2012). The presence of such common network-level perturbations explains why lesions in different locations in the brain produce remarkably similar behavioral deficits in different patients (Corbetta et al., 2018).

FC alterations suggest that behavioral deficits are due to the perturbation of inter-areal information flow. However, FC analyses cannot reveal the directionality or time scale of the information flow, leaving several questions open: i) is the stroke-related decrease of interhemispheric FC associated with a symmetric or asymmetric decrease in information flow between the damaged and non-damaged hemisphere? ii) is the increase of between-network intra-hemispheric FC paralleled by a change in intra-hemispheric information flow? iii) to which extent do changes in network-level information flows predict cognitive deficits? To address these questions, we performed covariance-based Granger Causality (GC) analyses (Brovelli et al., 2015) of resting-state fMRI data collected from stroke patients in the sub-acute phase (two weeks after stroke onset). Data were provided by the Washington university stroke database (Corbetta et al., 2015), and included structural lesions, resting-state fMRI, and neuropsychological scores for a large cohort of first-time stroke patients and age-matched control subjects. Analyses revealed that inter-hemispheric information transfer was significantly decreased in stroke patients with respect to healthy controls. In addition, pronounced inter-hemispheric imbalances in information transfer were observed in patients. Both anomalies were more prominent in resting-state networks related to attention and language, and they paralleled deficits in several behavioral domains.

## MATERIALS & METHODS

### Brain imaging and behavioral measurements

Details about participants, neuroimaging data acquisition and preprocessing, and brain lesion identification can be found in previous publications on the same data set (Corbetta et al., 2015, Siegel et al., 2016). Therefore, here we report only key information allowing for a self-contained reading of the paper.

#### Subject Enrollment and Retention

Participants (n = 172) were prospectively recruited. First-time stroke patients with clinical evidence of motor, language, attention, visual, or memory deficits based on neurological examination were included. One hundred and thirty-two patients met all inclusion criteria (for details see Corbetta et al., 2015) and completed the entire subacute protocol (mean age 52.8 years with range 22-77, 119 right-handed, 63 females, 64 right hemisphere). Patients were excluded from analysis for poor quality imaging data (n = 5), fewer than 400 frames remaining after motion scrubbing (n=8), or excessive hemodynamic lags (see below, n = 6) leaving 113 subjects in the final analysis. Demographically matched controls (n = 31) were recruited and underwent the same behavioral and imaging exams (mean age 55.7 years, SD = 11.5, range 21-83) in two separate scanning sessions (time point 1 and time point 2). Controls were matched to the study population in age, gender, handedness, and level of education. Controls were excluded based on a low number of frames after motion scrubbing (n = 4 at time point 1, n = 6 at time point 2), leaving 27 controls at time point 1 and 25 controls at time point 2.

#### Neuropsychological evaluation

Participants underwent a behavioral battery devised to assess motor, language, attention, memory, and visual function following each scanning session (details can be found in in Siegel et al., 2016). As described in Corbetta et al. 2015, principal components analysis was performed on all tests within a behavioral domain to produce a single score that predicted the highest percentage of variance across tasks. The left/right ‘Motor’ scores described left/right body motor performance that correlated across shoulder flexion, wrist extension/flexion, ankle flexion, hand dynamometer, nine-hole peg, action research arm test, timed walk, functional independence measure, and the lower extremity motricity index. The ‘Visual Field Attention’ score described contra-lesional attention biases in Posner, Mesulam, and behavioral inattention center-of-cancellation tasks. The ‘Sustained Attention’ score loaded on non-spatial measures of overall performance, reaction time, and accuracy on the same tests. The ‘Shifting Attention’ score loaded on tests indexing attention shifts, e.g. the difference in response times for attended *versus* unattended targets. The ‘Spatial Memory’ score loaded on the Brief Visuospatial Memory Test and spatial span. The ‘Verbal Memory’ score loaded on the Hopkins Verbal Learning Test. The ‘Language’ score loaded on tests devised to assess language comprehension (complex ideational material, commands, reading comprehension) and production (Boston naming, oral reading). The score of each of the seven factors for each patient was normalized using the mean and standard deviation of the corresponding factor scores in age-matched controls.

#### Brain imaging acquisition

Patients were scanned two weeks (mean = 13.4 days, SD=4.8 days) after stroke onset. Controls were scanned twice at an interval of 3-months. All imaging was performed using a Siemens 3T Tim-Trio scanner at the Washington University School of Medicine (WUSM) and a standard 12-channel head coil. The MRI protocol included structural, functional, pulsed arterial spin labeling (PASL), and diffusion tensor scans. Structural scans included: i) a sagittal T1-weighted MP-RAGE (TR = 1950 msec, TE = 2.26 msec, flip angle=90°, voxel size=1.0×1.0×1.0 mm); ii) a transverse T2-weighted turbo spin-echo (TR = 2500 msec, TE=435msec, voxel-size=1.0×1.0×1.0mm); and iii) sagittal FLAIR (fluid attenuated inversion recovery) with TR = 7500 msec, TE = 326 msec and voxel-size=1.5×1.5×1.5mm. Resting-state functional scans were acquired with a gradient echo EPI sequence (TR = 2000 msec, TE = 27 msec, 32 contiguous 4 mm slices, 4×4mm in-plane resolution) during which participants were instructed to fixate a small white cross centered on a screen with a black background in a low luminance environment. Six to eight resting state fMRI runs, each including 128 volumes (for a total of 30 minutes) were acquired. A camera fixated on the eyes was used to determine when a subject’s eyes were open or closed during scans. Patients had eyes open on 65.6±31.9% of frames and controls had eyes open on 76.8±30.2% of frames (t (114) = −1.7, p = 0.091).

#### Brain lesion masking

Lesions were manually segmented on individual structural MRI images (T1-weighted MP-RAGE, T2-weighted spin echo images, and FLAIR images obtained from 1 to 3 weeks post-stroke) using the Analyze biomedical imaging software system (www.mayo.edu; Robb and Hanson, 1991). Two board-certified neurologists (Dr. Maurizio Corbetta and Dr. Alexandre Carter) reviewed all segmentations. In hemorrhagic strokes, edema was included in the lesion. A neurologist (MC) reviewed all segmentations a second time paying special attention to the borders of the lesions and degree of white matter disease. Atlas-registered segmented lesions ranged from 0.02 cm^3^ to 82.97 cm^3^ with a mean of 10.15 cm^3^ (SD = 13.94 cm^3^). Lesions were summed to display the number of patients with structural damage for each voxel.

#### fMRI data preprocessing

Preprocessing of fMRI data included: i) compensation for asynchronous slice acquisition using sinc interpolation; ii) elimination of odd/even slice intensity differences resulting from interleaved acquisition; iii) whole brain intensity normalization to achieve a mode value of 1000; iv) removal of distortion using synthetic field map estimation and spatial realignment within and across fMRI runs; v) resampling to 3mm cubic voxels in atlas space including realignment and atlas transformation in one resampling step. Cross-modal (e.g., T2-weighted to T1-weighted) image registration was accomplished by aligning image gradients. Cross-modal image registration in patients was checked by comparing the optimized voxel similarity measure to the 97.5 percentile obtained in the control group. In some cases, structural images were substituted across sessions to improve the quality of registration. Following cross-modal registration, data were passed through three additional preprocessing steps. First, tissue-based regressors were computed based on FreeSurfer segmentation (Fischl, Sereno, Tootell, & Dale, 1999). The following sources of spurious variance were removed by regression: i) six parameters obtained by rigid body correction of head motion; ii) the signal averaged over the whole brain; iii) signal from ventricles and CSF; iv) signal from white matter. For Undirected Functional Connectivity (UFC) computations, we additionally regressed v) the average signal for gray matter. This step, commonly called global signal regression (GSR) was not applied for Granger causality (GC) computations. The rationale for this choice was to avoid any potential suppression of highly variable signals (Nalci et al., 2019) and distortion of information flow estimates using GC. Second, we performed temporal filtering retaining frequencies in the 0.009–0.08 Hz band. Third, we applied frame censoring meaning that the first four frames of each BOLD run were excluded. Frame censoring was implemented using frame wise displacement (Power et al., 2014) with a threshold of 1mm. This frame-censoring criterion was uniformly applied to all R-fMRI data (patients and controls).

#### Cortical surface processing

Surface generation and processing of functional data followed procedures similar to Glasser et al. (Glasser et al., 2013), with additional consideration for cortical segmentation in stroke patients. First, anatomical surfaces were generated for each subject’s T1MRI using FreeSurfer automated segmentation (Fischl et al., 1999). This included brain extraction, segmentation, generation of white matter and pial surface, inflation of the surfaces to a sphere, and surface shape-based spherical registration to the subject’s “native” surface to the fs average surface. Segmentations were manually checked for accuracy. For patients in whom the stroke disrupted automated segmentation, or registration, values within lesioned voxels were filled with normal atlas values prior to segmentation, and then masked immediately after (7 patients). The left and right hemispheres were then resampled to 164,000 vertices and registered to each other (Van Essen et al., 2001), and finally down-sampled to 10,242 vertices each (a combined total of 18,722 vertices after exclusion of the medial wall) for projection of functional data. Following preprocessing, BOLD data were sampled to each subject’s individual surface (between white matter and pial surface) using a ribbon-constrained sampling available in Connectome Workbench (Marcus et al., 2013). Voxels with a high coefficient of variation (0.5 standard deviations above the mean coefficient of variation of all voxels in a 5 mm sigma Gaussian neighborhood) were excluded from volume to surface mapping (Glasser et al., 2013). Time courses were then smoothed along the 10,242 vertex surface using a 3mm FWHM Gaussian kernel. All brain surface visualizations were generated using Connectome Workbench (Marcus et al., 2013).

#### Brain parcellation scheme

We used a cortical surface parcellation generated by Gordon & Laumann and colleagues (Gordon et al., 2016). The parcellation is based on R-fMRI boundary mapping and achieves full cortical coverage and optimal region homogeneity. The parcellation includes 324 regions of interest (159 left hemisphere, 165 right hemisphere). Note that the original parcellation includes 333 regions, while here all regions less than 20 vertices (approximately 50 mm^2^) were excluded. This cutoff was arbitrarily chosen based on the assumption that parcels below this size would have unreliable signal given 4 mm sampling of our functional data. Notably, the parcellation was generated on 120 young adults aged 18-33 and is applied here to adults aged 21-83. To generate parcel-wise connectivity matrices, time courses of all vertices within a parcel were averaged. For each ROI, we defined its center-of-mass coordinates 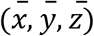 as the average of the (*x,y,z*) coordinates of all vertices in the ROI. For each ROI, identified the homologous regions as the ROI in having the lowest distance from 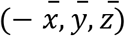(i.e., the ROI closest to be symmetrically located in the opposite hemisphere).

In addition to the 324 cortical parcels, we also defined a set of 19 sub-cortical and cerebellar regions based on the FreeSurfer segmentation: for each hemisphere 9 regions consisting of cerebellum, thalamus, caudate, putamen, pallidum, hippocampus, amygdala, accumbens and ventral dorsal caudate, plus brainstem (Fischl et al., 2002).

### Granger causality analysis and inter-areal information transfer

#### Granger Causality (GC) framework

One of the most successful data-driven methods to quantify the degree of communication from statistical dependencies between neural signals is based on the Wiener-Granger causality principle (Granger, 1980; Brovelli et al., 2004; Ding et al., 2006; Bressler and Seth, 2011; Seth et al., 2015). In order to describe the Granger causality framework, let us consider two (discrete) time series *X* = {*X_t_*},*Y* = {*Y_t_*} representing the activity of two subsystems sampled at discrete times *t={1,2,3,…,n}*, where we assume that times are measured in units of the sampling time TR. Standard undirected functional connectivity (UFC) is classically computed as the Pearson’s correlation, defined as 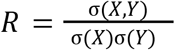 where σ(*X*) and *σ*(*Y*) are respectively the standard deviations of *X_t_* and *Y_t_* and σ^2^ (*X, Y*) is their covariance within the selected time window. The UFC only considers dependencies between *X_t_* and *Y_t_* for the same *t*. Information-theoretically, this type of dependency is quantified by the mutual information *I*(*X_t_*; *Y_t_*), which is a simple function of *R* for Gaussian data, *I*(*X_t_*; *Y_t_*) =- 1/2*log*(1 – *R*^2^). The UFC is insensitive to the temporal structure of correlation between *X* and *Y,* since it is invariant under permutation of *t*. On the other hand, the framework based on Granger causality (Granger 1963; 1980) and further developed by Geweke (Geweke, 1982) considers dependencies between two time series and their “lagged” versions with different lags 1,2,…. Let us assume that *L* is the maximum lag at which dependence is observed: in other words, *X_t_, Y_t_* are not dependent on *X*_τ_, *Y*_τ_- for *τ* < *t* < *L*, i.e., he values of *X* and *Y* occurring before a time *t* − *L* in the past. One can thus restrict attention to dependencies between *X_t_, Y_t_* and the *L* preceding values of the time series,

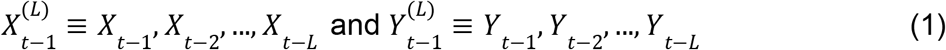

Interdependencies between the two time series reflected into the fact that the time series of *Y* contains information about *X_t_*, and vice versa. The total amount of information about *X_t_* contained in *Y* can be quantified by the mutual information between *X_t_* and *Y*(present and past): 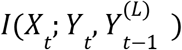. By virtue of the identity *I*(*A*;*BC*) = *I*(*A*;*B*) + *I*(*A*;*C*|*B*), this quantity can be decomposed into an “instantaneous” and a “lagged” term:

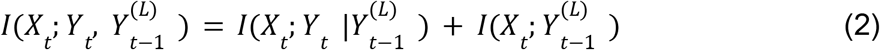

The conditioning on the instantaneous term implies that 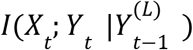 measures information about *X_t_* contained exclusively in *Y_t_* (and not already contained in 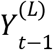). The basic idea of Granger causality is that *Y* contains exclusive information about *X_t_* which is not already present in the time series of *X*, i.e., in 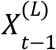. To obtain the “exclusive” information about *X_t_* contained in *Y* one should further condition over 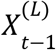:

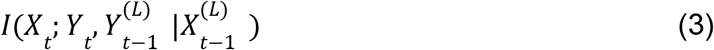

and again obtain an “instantaneous” and a “lagged” term:

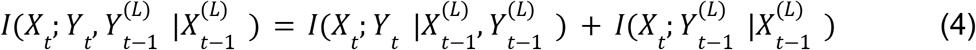

The first term is called instantaneous causality (IC) and usually indicated by *F_X·Y_*

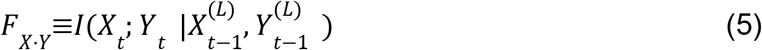

The second term is called directed causality (DC) from *Y* to *X* and usually indicated by *F_Y→X_*

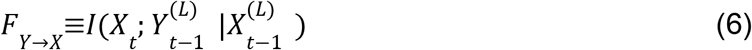

Symmetrically, the exclusive information about *Y_t_* contained in *X* is measured by

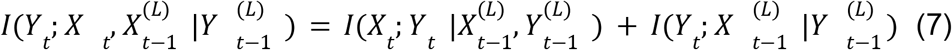

The first term coincides with *F_X·Y_* and the second one is the directed causality from *X* to *Y*,

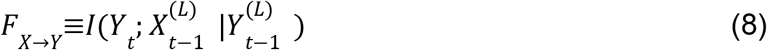

The measures *F_X→Y_*, *F_Y→X_* and *F_X·Y_* were proposed by Geweke (Geweke, 1982), who also defined the total interdependence between *X* and *Y* as the sum of all the three terms,

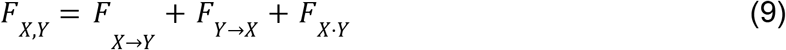

This is the “new” dependency between *X* and *Y* “created” at each time *t*, indeed

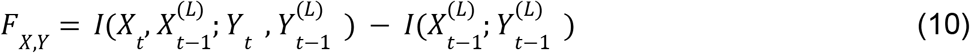

Thus, in the GC framework, the total interdependence between two signals can be split into three terms: two directed Granger causality (DC) terms and an instantaneous Granger causality (IC) term. The DC terms (*F_X→Y_*, *F_Y→X_*) represent a directed flow of information from *X* to *Y* or vice versa, occurring over a timescale greater than 1 TR (fig. 1b). The IC term (*F_X·Y_*) represents information shared between *X* and *Y* “instantaneously”, i.e., in less than one TR, and it accounts for direct communication, or unconsidered influences that may originate from common (e.g., subcortical) sources, occurring in less then one TR (fig. 1a).

**Figure 1.**
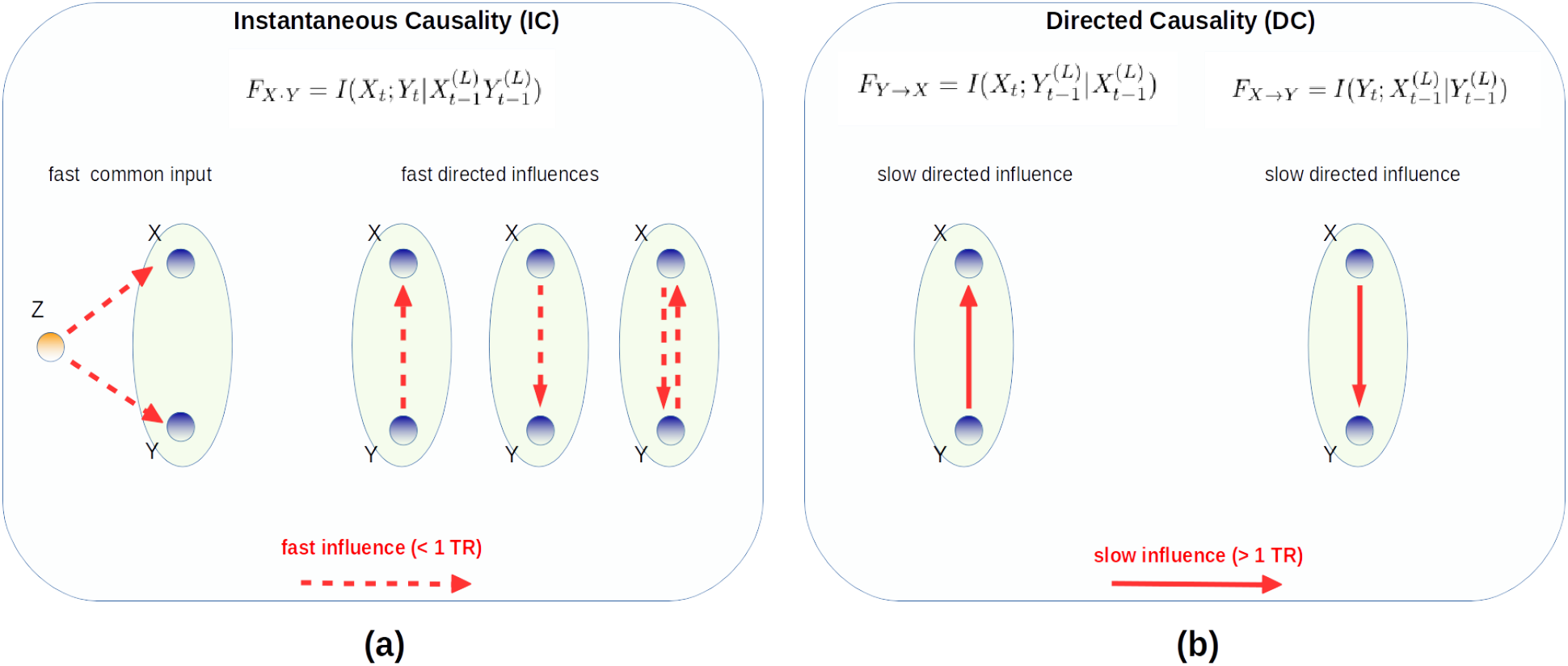
Interpretation of instantaneous and directed Granger causality in terms of information flows between two areas.

Granger causality measures can therefore be formulated in completely information-theoretical terms (Barnett et al., 2009; Marko, 1973; Rissanen and Wax, 1987; Schreiber, 2000). Information-theoretic measures based on the Wiener-Granger principle, such as Transfer Entropy (Schreiber, 2000) and Directed Information (Massey, 1990), represent the most general measures of Wiener-Granger causality and capture any (linear and nonlinear) time-lagged conditional dependence between neural signals (Besserve et al., 2015; Vicente et al., 2011).

#### Covariance-based Granger Causality

The GC measures *F_X→Y_, F_Y→X_* and *F_X·Y_* capture statistical relations among the values of *X,Y* in a time window of length *L* + ļincluding the “present” values *X_t_, Y_t_* and the “past” values 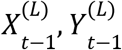. Together, 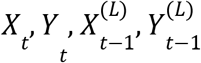 define a vector of length 2*L* + 2 values. The GC measures can be ultimately expressed in terms of Shannon entropies involving the (2*L* + 2-variate) probability distribution 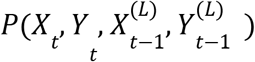 and some of its marginals:

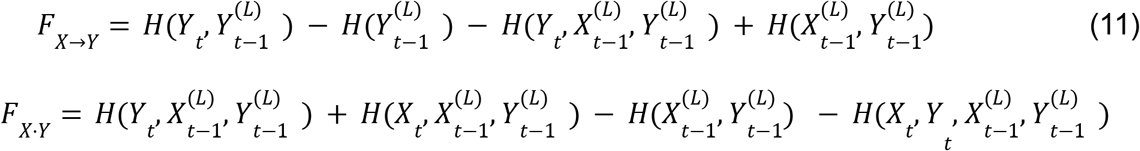

Assuming the distribution 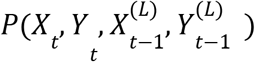 to be stationary, the classical method to compute entropies is the “binning method”. One considers *T* running windows of length *L* + 1, and for each window extracts the vector 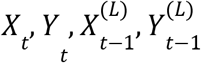, thus obtaining *T* samples of 2*L* + 2. Binning each univariate variable and collecting the bin counts, the joint probability distribution 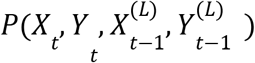 is approximated by (multidimensional) histogram (Beirlant et al., 1997; Treves and Panzeri, 1995). If *n* bins are used for each univariate variable, the total number of multidimensional bins is *n*^2(*L*+1)^. As a rule of thumb, to get at least a rough estimate of the bin counts one needs at least as many samples as bins, so *T*≥*n*^2(*L*+1)^ points. Since *L*≥1, this requires a large sample *T*≥*n*^4^ for estimation. In order to make the estimation feasible on short time windows, a common solution is to approximate the distribution with the first term of the Gram-Charlier expansion, i.e., by a Gaussian distribution with the same second order moments (covariance matrix) as the given distribution. This approximation amounts to keeping only second order statistics, and neglecting higher-order terms, and is relatively accurate for fMRI data (Hlinka et al., 2011). In this approximation, the distribution 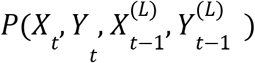 and its marginals are effectively replaced by the covariance matrix 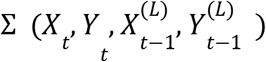 and its submatrices. Estimating Σ requires only to estimate (2*L* + 2)^2^ parameters corresponding to the second moments of the distribution. Furthemore, entropies can be simply computed with the formula

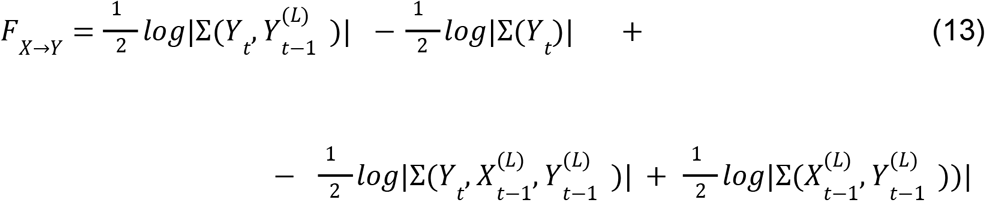

where *Σ*(*A*) is the covariance matrix of *A, n_A_* the dimension of *A*,> and |·| is the determinant. In this *covariance-based* approximation, GC measures are expressed in terms of determinants of submatrices of the covariance matrix of the data (Brovelli et al., 2015). For instance,

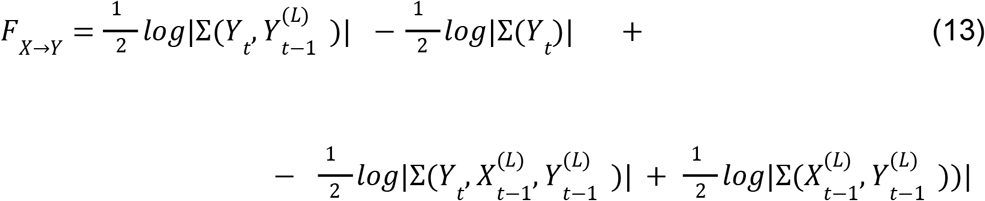

The covariance-based GC estimation is equivalent to the parametric estimation of GC from an autoregressive model with Gaussian innovations, i.e., the traditional way to estimate Granger causality.

#### Gaussian-copula-based estimation of GC

The GC measures *F_X→Y_*, *F_Y→X_* and *F_X·Y_* can all be written as appropriate sums of mutual information (MI) terms. For instance,

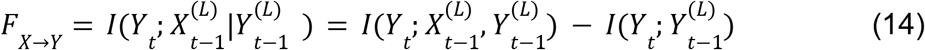

The MI is invariant under monotonic transformations of the marginals, and this fact can be exploited to relax the assumption of Gaussianity. In particular, one can replace the assumption of Gaussianity with the weaker assumption of a *Gaussian copula* (Ince et al., 2017): that the joint distribution of the variables can be rendered Gaussian by means of local transformations on the marginals. Formally, consider the transformations

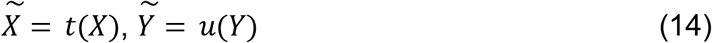

where *t, u* are monotonic functions. The Gaussian copula assumption is equivalent to the existence of *t, u* such that 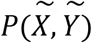 is Gaussian. While Gaussianity imposes a linear dependence between variables, the Gaussian copula assumption allows for more general monotonic dependence (Ince et al., 2017). Under the Gaussian copula assumption, there is a simple way to compute the MI. Since the MI is invariant under monotonic transformations of the marginals, 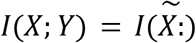, and since there exist 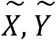 such that 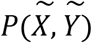 is Gaussian, it is sufficient to find 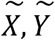 and compute 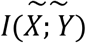 with the covariance-based formula (12), which is exact for 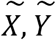.

Finding 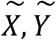 is easy. Consider two random variables *X,Y* with joint cumulative distribution function (CDF) *H*(*X, Y*) and marginal CDFs *F*(*X*), *G*(*X*). Consider

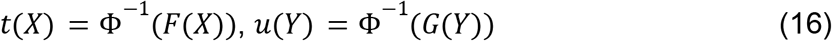

where Φ is the CDF of a standard normal variable. One can immediately show that 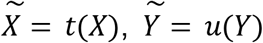 are standard normal variables, i.e., 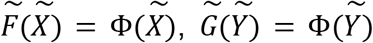. Applying the covariance-based approach to the transformed variables, one has 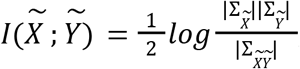, and by virtue of the invariance 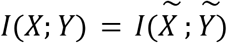.

The Gaussian copula evaluation of the MI is not exact. In general, even though the marginal distribution of 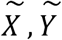 are Gaussian, the joint distribution is not perfectly a bivariate Gaussian. This is true only if the *copula,* i.e., the part of the distribution specifying the dependence between the two variables, is Gaussian (see Ince et al. 2017 for a definition of copula and discussion of this point). However, for many distributions the Gaussian copula assumption is approximately met. In summary, one obtains the following algorithm to compute the MI: i) given samples {*X_t_,Y_t_*}, approximate *F*(*X*), *G*(*Y*) with the empirical CDFs *F*(*X_i_*) = *rank*(*X_i_*)/*N*, *G*(*Y_i_*) = *rank*(*Y_i_*)/*N* and compute *t*(*X*) = Φ^-1^ (*rank*(*X_i_*)/*N*), *u*(*Y_i_*) = Φ^-1^ (*rank*(*Y_i_*)/*N*). *t*(*X_i_*) and *u*(*Y_i_*) are normally distributed and have the same MI as *X_i_,Y_i_*. ii) compute the MI from the samples {*t*(*X_i_*),*u*(*Y_i_*)} with the covariance-based method. In our work, we have computed all GC measures by expressing them in terms of sums of MIs and then applying the Gaussian-copula-based estimation to each term in the sum.

Overall, the first advantage of bivariate gaussian-copula and covariance-based GC is its applicability to large networks of nodes, such as the 343 nodes used in this study. Most other methods to infer directed GC (such as multivariate Granger causality) are more accurate in inferring *direct* (i.e., non-mediated) influences, but usually applicable to only smaller networks (of the order of 100 nodes) (Tang et al., 2012; Stramaglia et al., 2016). The use of covariance-based GC does not require to select a specific sub-network of nodes, or to average BOLD signals over large regions (which would imply a considerable signal loss, due to potential inhomogeneities). A second advantage is its estimability from short signals. This property allows us to estimate GC from BOLD time series of 400-800 time points, i.e., the time series of single subjects. Thus, we do not need to concatenate several subjects to perform the estimation, and we can obtain individual estimates. These two key properties enable a direct comparison of Granger causality results with previous FC studies.

#### Choice of the appropriate lag (L)

GC measures *F_X→Y_, F_Y→X_* and *F_X·Y_* depend on the maximum lag *L* used. Intuitively, *L* should be as large as to include all points in the past that have correlations with *X_t_, Y_t_*. This would roughly correspond to the autocorrelation decay time of the time series. For rs-fMRI, this is of the order of 10s (5 TR). More rigorously, we tested how many points in the past significantly contribute to predicting the present values of the time series by assuming a specific form for the process generating the data. If we assume that the data are generated by a Gaussian vector autoregressive (VAR) process of length *L*

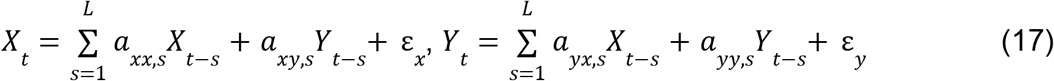

where ε_*x*_, ε_*y*_ are (possibly correlated) Gaussian innovations. This can be rewritten as

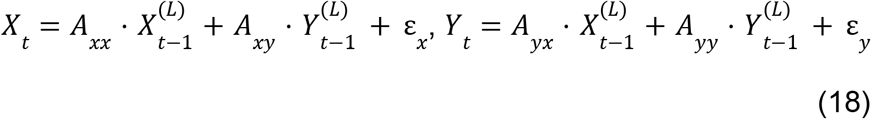

with *L*×1 vectors *A_xx_, A_xy_, A_yx_, A_yy_*. In this setting *X_t_, Y_t_* are Gaussian with means 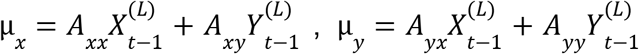 and covariance *σ* given by the 2×2 covariance matrix of ε_*x*_, ε_*y*_. Using least squares estimation, one obtains the best fit for the parameters

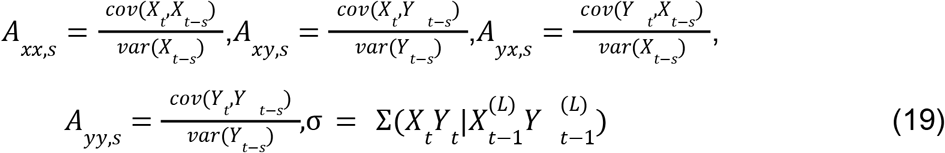

where Σ(*A*|*B*) = Σ(*A*) – Σ(*X Y*)Σ^-1^(*B*)Σ(*A,B*)^*T*^ is the partial covariance matrix of *A* given *B*, with 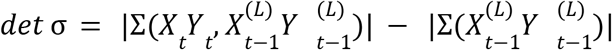.

An appropriate choice of *L* is given by model comparison, i.e., by fitting models with different *L* to the data and selecting the model yielding the “best fit”. Since the models for increasing *L* are nested (models with higher *L* include models with lower *L* as a special case), the likelihood of the models increases monotonically as a function of *L.* To avoid overfitting, the customary procedure with nested models is to select the proper *L* by a model-comparison criterion penalizing models with a larger number of parameters. Here, we used the common BIC criterion (McQuarrie and Tsai, 1998) that assesses the fitness of each model as *B* = 2*log*Λ – *d log*(*n*) where Λ is the log-likelihood, and *d log*(*n*) is a term penalizing models with a large number of parameters *d*. The log-likelihood of the model (17) is 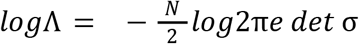 *det σ* which gives the BIC value

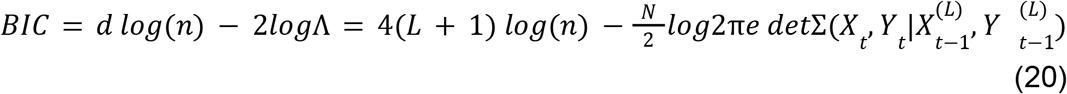

The best-fitting model is the one maximising *BIC.* We computed the value of *BIC* for all pairs of regions-of-interest (ROIs) and all subjects (patients and controls) as a function of *L.* We found that on average BIC increases up to *L*≈5, and then remains relatively stable. Note that since TR= 2s, *L* = 5 corresponds to the expected autocorrelation length of the signal. On the basis of these results, we chose to fix *L* = 5 in our analyses.

#### Relation between linear correlation and covariance-based Granger causality measures

Standard undirected functional connectivity (UFC) is classically computed as the Pearson’s correlation, defined as 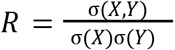 where σ(*X*), σ(*Y*) are the standard deviations of *X_t_* and *Y_t_* and σ (*X, Y*) is their covariance within the selected time window. The UFC only considers dependencies between *X_t_* and *Y_t_* for the same *t*. UFC and covariance-based Granger causality measures share common properties. Linear correlation and total interdependence are undirected measures quantifying static and dynamic dependencies, respectively. Although these measures are not related by a mathematical decomposition, there is a strong relationship between the existence of both types of dependencies. A lack of total interdependence implies a lack of linear correlation; and, if we assume that the future of *X* and *Y* causally depends on their own past, respectively, the opposite relation is also true. This occurs because linear correlation is related to the covariance-based approximation of the mutual information, *I*(*X_t_*; *Y_t_*) =< 1/2*log*(1 < *R*), and because conditioning on the past cannot create new dependencies (Chicharro and Panzeri, 2014). It is also clear the directed and instantaneous Granger measures are smaller than the total interdependence. Thus, null total interdependence implies the absence of Granger causality measures because they constitute non-negative contributions to the total interdependence. In other words, Granger causality is present if, and only if, both linear correlation and total Granger interdependencies are not zero.

#### The FC and GC quality-based exclusion criteria

In order to ensure good-quality FC and GC estimates, we excluded from analysis all subjects with less than 400 usable frames after motion scrubbing. Furthermore, for each subject, we computed a lag between homologous ROIs as in (Siegel et al., 2016B). In brief, for any integer lag *l* =< 4, < 3,…, 3,4 we computed the lagged cross-correlation *C_l_* = < *X_t_Y*_*t*+1_ > between the BOLD signals *X,Y* of the homologous ROIs; the homotopic lag between the ROIs was identified by finding *l*_0_ = *argmin*(*C_j_*), performing a parabolic interpolation on 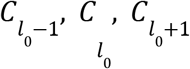, and computing the minimum of the parabola. An average homotopic lag between the left and right hemisphere was computed by averaging over all homptipic lags between left ROIs and the homologous right ROIs. Anomalously large homotopic lags are a likely indication of the presence of lags of hemodynamic origin, due to disruption of the standard hemodynamic response in the vicinity of the lesion. Therefore, we excluded from analysis all subjects with severe homotopic lags (greater than 1s inter-hemispheric difference). After motion and lag exclusion, 113 patients were included at two weeks, 27 controls at time point one, and 25 at time point two.

## RESULTS

We analyzed resting-state fMRI data recorded from acute stroke patients (n=113) and healthy participants(n=26). Our analysis tested the hypothesis that post-stroke FC alterations are tightly intertwined with information flow deficits occurring both inter-hemispherically and intra-hemispherically. To address this issue, we performed covariance-based Granger causality (GC) analyses (Brovelli et al., 2015) of resting-state fMRI data and compared inter-areal information flow analyses with standard FC approaches. The comparison was performed by means of the notion of total interdependence between signals (Geweke et al., 1982). In the GC framework, the total interdependence (TI) between two signals can be split into three terms: two directed Granger causality (DC) terms and an instantaneous Granger causality (IC) term. The DC terms (*F_X→Y_, F_Y→X_*) represent a directed flow of information from *X* to *Y* or vice versa, occurring over a timescale greater than 1TR (fig. 1b). The IC term (*F_X·Y_*) represents information shared between *X* and *Y* “instantaneously”, i.e., in less than one TR, and it accounts for rapid direct communication, or unconsidered influences that may originate from common (e.g., subcortical) sources (fig. 1a). Functional MRI data were computed for 324 parcels of the Gordon-Laumann cortical parcellation (Gordon et al., 2016) and 19 sub-cortical and cerebellar parcels from the FreeSurfer atlas (Fischl et al., 2002). For each subject and for each pair of parcels, or regions-of-interest (ROIs), we evaluated the undirected functional connectivity (UFC, z-transformed Pearson correlation), the instantaneous Granger causality (IC) and the directed Granger causality in both directions (DC). For healthy controls, the DC, IC and UFC matrices obtained in two independent sessions were averaged.

### Consistency of UFC and GC measures across fMRI sessions

We first tested the reliability of our results by verifying the consistency of the UFC and GC measures obtained for control participants in two separate sessions (Fig. 2a). Consistency was defined as the Pearson correlation between the (upper-triangular parts of the) corresponding matrices in the two sessions. The UFC matrices were highly consistent (r=0.65±0.03, average and standard error over subjects). The same result was obtained for IC matrices (r=0.73±0.02). The DC matrices, instead, were much less consistent (r=0.22±0.02). We obtained reduced network-wise (28×28) matrices by averaging over ROIs in the same network and hemisphere. We considered thirteen cortical resting-state networks as in (Gordon et al., 2016), plus subcortical ROIs. Consistency improved for UFC (r= 0.80±0.03), IC (r=0.87±0.03), and DC (r=0.41±0.03). The UFC and IC results are thus reliable at the single-subject level, especially if network-averaged results are considered. As for the DC, due to the poor level of consistency obtained in the full (343×343) DC matrix, we cannot expect reliable results at the level of single subject, single ROI. Also at the network level individual results are not completely reliable. To assess the reliability of group results, we computed the consistency of group-averaged FC matrices for random groups of *n* participants(Fig. 2b). The group consistency is significantly stronger than the individual consistency. When considering groups of 5 subjects, the DC consistency rises to 0.4 (0.7 for network-wise matrices), and for 10 subjects it rises to 0.5 (0.8 for network-wise matrices). This result implies that while individual DC results are affected by a very large noise, DC results at the group-level are reliable. In summary, UFC and IC matrices were highly consistent both at the individual and group level, while DC matrices were consistent only at the group level.

**Figure 2:**
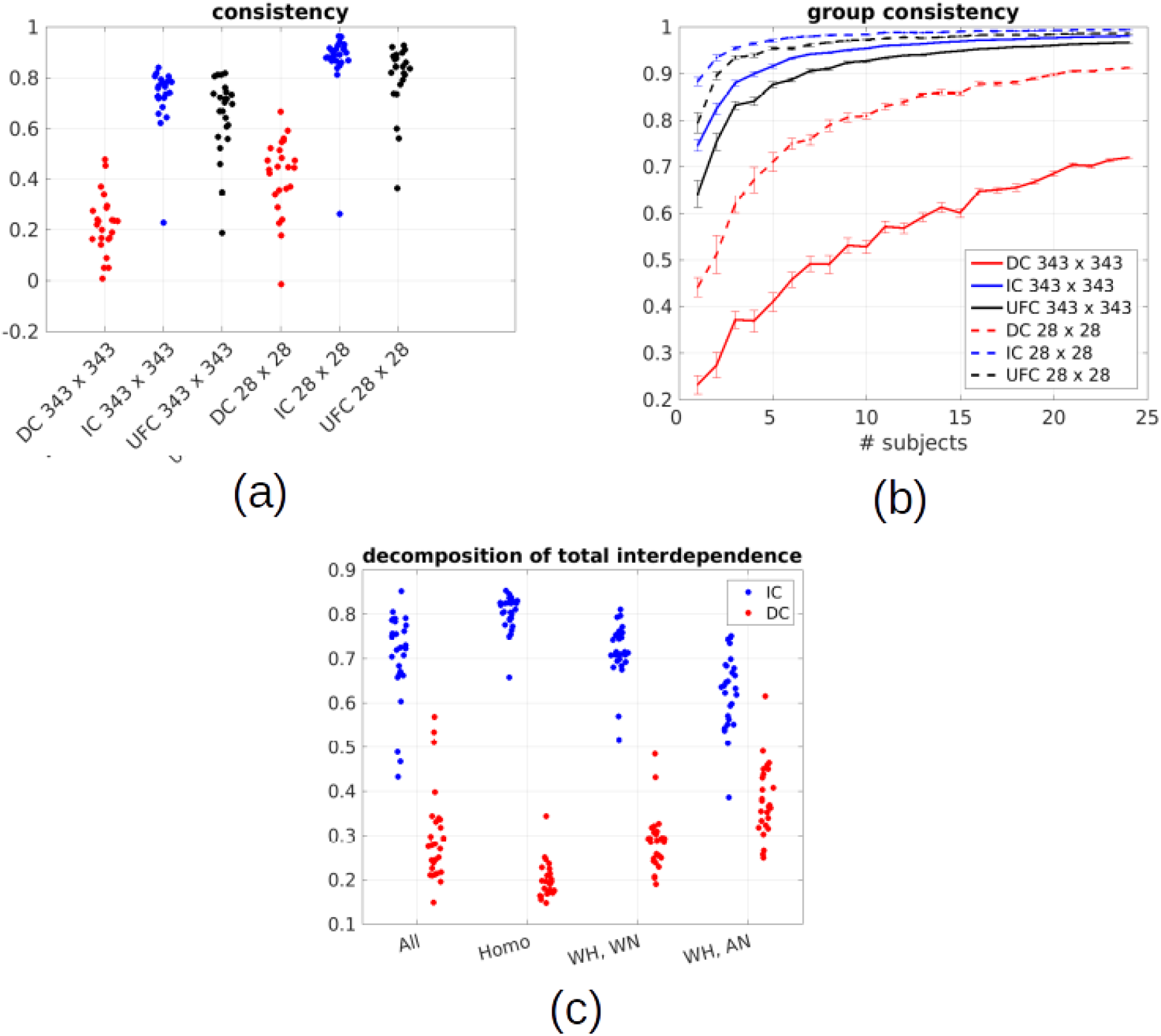
(a-b) Consistency of FC measures in two separate sessions. We computed 343×343 DC, IC and UFC matrices for the two separate sessions of control subjects. We assessed consistency between the FC results in the two sessions as the Pearson correlation between the (upper-triangular part of the) corresponding matrices in the two sessions. We also evaluated consistency between the 28×28 DC, IC and UFC matrices obtained by averaging over all pairs of ROIs belonging to the same RSN (13 cortical resting state systems + subcortical ROIs). In panel (a) we show the distribution of consistency for the individual results of each subject. In panel (b), we assess consistency of group averages. For each *n*, we randomly select *n* subjects and average the FC matrices over subjects. We then show the average (over random choices of n subjects) consistency of the group-averaged matrices as a function of *n.* (c) average fraction of the total interdependence (TI) accounted for by the instantaneous causality (IC) and directed causality (DC), for each subject and different classes of links: all links (all), homotopic links (homo), within-hemisphere within-RSN links (WH, WN), within-hemisphere across-RSN links (WH, AN) links. It is apparent that IC accounts for a large fraction of the TI, especially for homotopic links (~80%).

To better understand the relation between the degree of consistency across sessions and the variance explained by each metric, we computed the fraction of TI due to the IC and DC. In fig. 2c, we show the proportion over the TI averaged across links for each individual subject. Overall, he IC accounts for a large fraction (mean: 70%, s.d.: 10%) of the TI. This fraction is even higher if we compute the mean over homotopic links, which are the strongest functional links, connecting homologous ROIs located symmetrically in opposite hemispheres (mean: 80%, s.d.: 4%). Another class of strong functional links is given by intra-hemispheric links within the same resting state network: also in this case, the largest fraction of the TI is due to the IC (mean: 71%, s.d.: 6%). If we consider weak functional links connecting different resting state networks, the fraction of TI due to the IC decreases, but remains well above 50% (mean: 61%, s.d. 8%). These results imply that most of the correlation between the time series of different ROIs is due to interactions occurring within the time resolution of fMIR (1TR=2s). This limits the detectability of directed interactions (DC), and consequently also the resolvability of directionality.

### Interhemispheric homotopic undirected functional connectivity and Granger causality analyses

Previous studies have shown that stroke patients present a reduced interhemispheric UFC with respect to healthy controls (Carter et al., 2010; Golestani et al., 2013; He et al., 2007; New et al., 2015; Park et al., 2011; Ramsey et al., 2016; Siegel et al., 2016; Tang et al., 2016). Such effect is strongest for interhemispheric homotopic connections, which link homologous ROIs located symmetrically in opposite hemispheres. We computed UFC, IC, and DC between pairs of homologous ROIs and compared healthy controls (C, n=26) with stroke patients (P, n=113) in the subacute phase. Furthermore, we subdivided the latter group into patients with lesions in the left hemisphere (LHP, n=60) and patients with lesions in the right hemisphere (RHP, n=53). LHP and RHP were kept separate as they were clinically different and presented dissimilarities in FC.

Global measures of homotopic connectivity were obtained by averaging the UFC, IC and DC over all pairs of homotopic links. Network-wise measures were obtained by averaging over pairs of homotopic ROIs belonging to the same resting state network (RSNs). We considered twelve RSNs as in (Gordon et al., 2016), in addition to subcortical regions, and left out networks with less than 5 nodes. We considered the following RSNs: the visual network (VIS), sensorimotor dorsal network (SMD), sensorimotor ventral network (SMV), auditory network (AUD), cingulo-opercular network (CON), ventral attention network (VAN), dorsal attention network (DAN), default mode network (DMN), fronto-parietal network (FPN), subcortical regions (SUB).

Figure 3a shows the distribution of the homotopic UFC, averaged over all homotopic pairs. We observed a significant difference in the UFC distribution between healthy controls, LH and RH patients. The UFC was significantly higher in controls than LH and RH patients, which in turn were not different (*one way ANOVA with group as factor: F(2,136)=6.6, p=0.002; post-hoc T-tests C vs LHP, T(84)=4.1, p=0.0001, C vs RHP T(77)=3.2, p=0.0001, LHP vs RHP T(111)=-0.9, p=0.37*). Figure 4d shows UFC distributions for each RSN. Controls had significantly higher UFC than LH patients in all networks except for the VAN, and significantly higher UFC than RH patients in all networks except for the VAN, DMN, FPN and SMV (*two-way repeated-measures ANOVA with group and network as factors; group F(2,1224) = 6.4, p=0.002; network: F(9,1224) = 126.7, p <10^-10^; interaction: F(18,1224) = 2.1, p= 0.005; post-hoc T-tests C vs LHP: p<0.05 in all networks except VAN, FDR-corrected for 10 comparisons; C vs RHP: p<0.05 in all networks except VAN, DMN, FPN and SMV, FDR-corrected for 10 comparisons*). In summary, stroke patients presented an overall decrease of homotopic UFC with respect to healthy subjects. The effect was strongest for homotopic connections in VIS, SMD, AUD, CON, DAN networks, and for subcortical regions. Analogous results were already obtained in previous analyses (Joshua Sarfaty Siegel et al., 2016).

**Figure 3:**
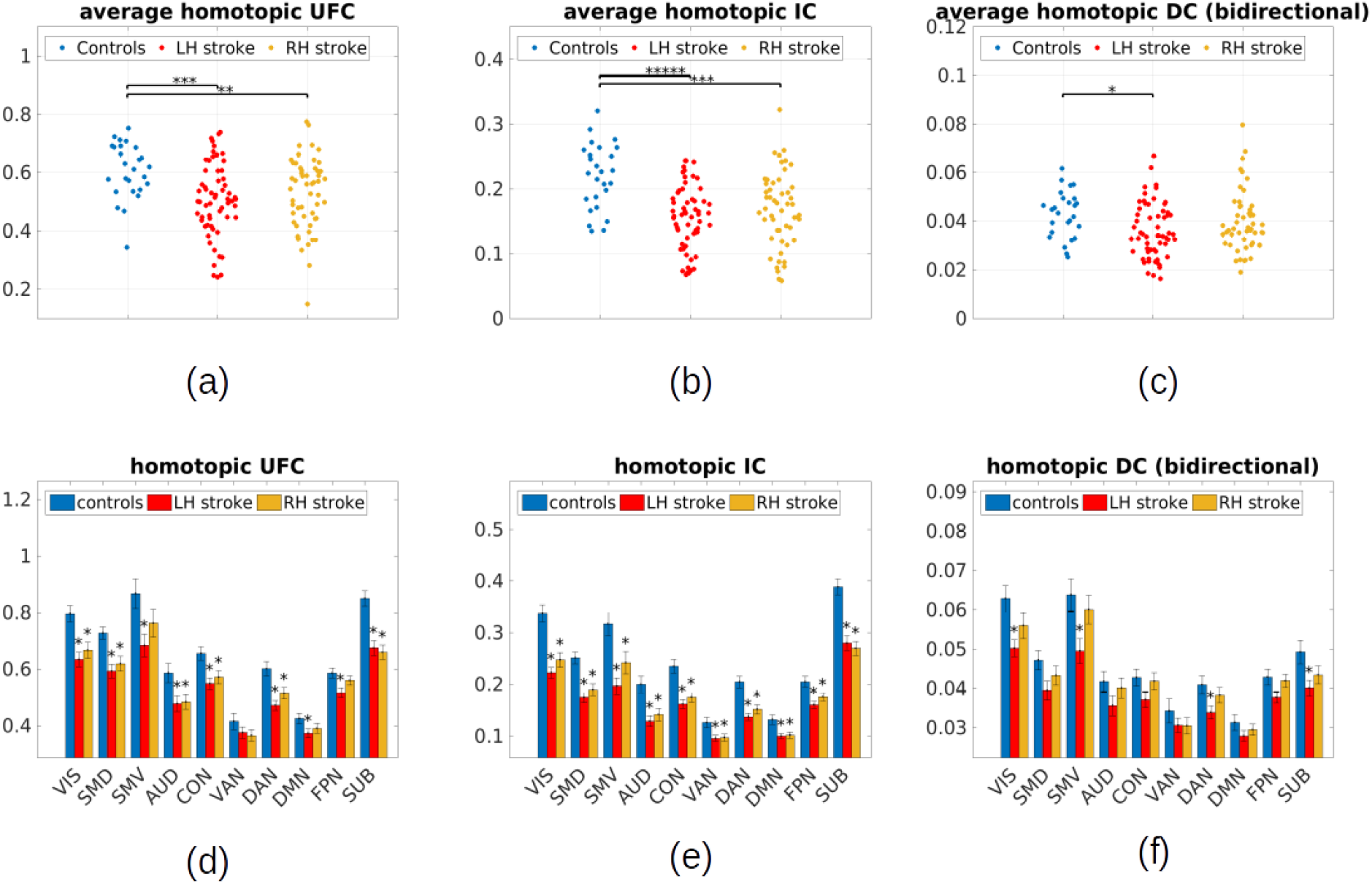
Average homotopic UFC, IC and DC in acute phase. For each subject, we averaged the UFC, the IC, and the bidirectional DC for homotopic pairs of ROIs. (a-c) show individual averages of homotopic UFC, IC, bidirectional DC (each dot represents one subject). At a group level, the average homotopic UFC and IC are higher for controls than both LH and RH patients. The bidirectional DC is higher for controls than LH patients. (d-f) show homotopic UFC, IC and bidirectional DC by resting-state network. Column heights are averages over subjects, error bars standard errors over subjects. At the group level, the UFC and IC for each network is higher in controls compared to patients. Control/patient differences are stronger for IC than UFC. The average bidirectional DC for each network is higher in controls compared to LH patients. Stars indicate networks for which comparison with controls (two-sample T-test, p<0.05 FDR corrected for 10 comparisons) is significant. VIS=visual, SMD=sensorimotor dorsal, SMV=sensorimotor ventral, AUD=auditory network, CON=cingulo-opercular network, VAN=ventral attention network, DAN=dorsal attention network, DMN=default mode network, FPN=fronto-parietal network SUB=subcortical nodes.

**Figure 4:**
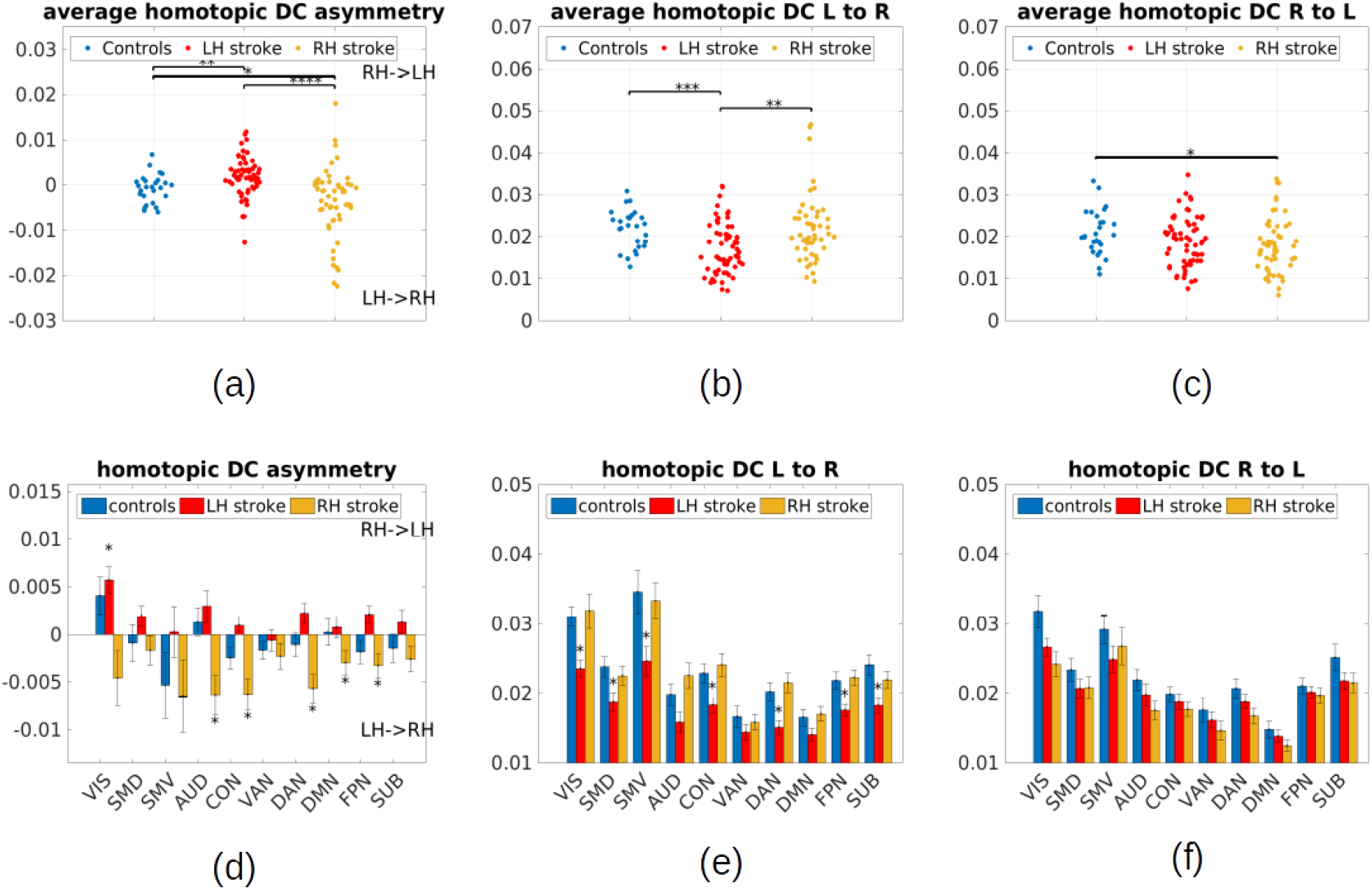
Direction of the average homotopic DC in acute phase. For each subject, we averaged the DC over homotopic pairs of ROIs, considering separately the DC from left regions to right regions, from right regions to left regions, and the difference, called homotopic DC asymmetry asymmetry. (a-c) we show individual averages of homotopic DC from left to right, right to left, and asymmetry (left to right - right to left). Each dot represents a subject. At a group level, for both LH and RH patients, the information flow (DC asymmetry) is in the direction of the lesioned hemisphere, i.e., the homotopic DC from the intact to the lesioned hemisphere is higher than vice versa. DC from left to right is reduced for all patients compared to controls, but much more strongly for LH patients. DC from right to left is reduced for RH patients compared to controls. Overall, DC from the lesioned to the intact hemisphere tends to be reduced in patients. (d) We show homotopic DC asymmetry (right to left - left to right) by resting-state network. Column heights are averages over subjects, error bars standard errors over subjects. For all networks, the information flow (DC asymmetry) is from right to left LH patients, and from left to right in RH patients. Stars represent networks for which comparison with 0 (one-sample T-test, p<0.05 FDR corrected for 10 comparisons) is significant. (e-f) We show homotopic DC from left to right and right to left. Column heights are averages over subjects, error bars standard errors over subjects. DC from left to right is significantly reduced in several networks for LH patients. Stars represent networks for which comparison with controls (two-sample T-test, p<0.05 FDR corrected for 10 comparisons) is significant. VIS=visual, SMD=sensorimotor dorsal, SMV=sensorimotor ventral, AUD=auditory network, CON=cing VIS=visual, SMD=sensorimotor dorsal, SMV=sensorimotor ventral, AUD=auditory network, CON=cingulo-opercular network, VAN=ventral attention network, DAN=dorsal attention network, DMN=default mode network, FPN=fronto-parietal network SUB=subcortical nodes.

The analysis of homotopic UFC in patients showed that the activity of homologous regions is less synchronized than in healthy controls, suggesting a reduced interaction between the hemispheres. Granger causality (GC) analyses were performed to characterize instantaneous (IC) and directional (DC) interactions between homologous regions. Figure 3b depicts the distribution of the instantaneous causality (IC), averaged over all homotopic pairs of ROIs. The IC was significantly higher in controls with respect to LH patients and RH patients, but did not differ between LH and RH patients (*one-way ANOVA with group as factor, F(2,136) = 14.0, p= 3·10^-6^; post-hoc t-tests C vs LHP, T(84) = 5.5, p=4·10^-6^, C vs RHP T(77) = 4.2, p=0.0001, LHP vs RHP T(111)=-1.1, p= 0.27*). Figure 3e shows results separately for each network. While controls had significantly higher IC than both LH and RH patients in all networks, the strongest differences were observed in the VIS, SMD, CON, DAN and subcortical regions (*two-way repeated-measures ANOVA with groups ad network as factors, group: F(2,1224)=13.5, p=4·10^-6^; network: F(2,1224)=180.3, p<10^-10^; interaction: F(18,1360)=4.2, p=1.0·10^-10^; post-hoc T tests C vs LHP, p < 0.05 in all networks, FDR-corrected for 10 comparisons; C vs RHP, p < 0.05 in all networks, FDR-corrected for 10 comparisons*). In summary, the IC results are in qualitative agreement with those of UFC, but the discrepancy between patients and control subjects is more pronounced (as mirrored in a larger group effect).

We then analyzed directional Granger causality measures between homologous regions. We first investigated whether the bidirectional information flow across hemispheres was different between patients and controls. To this aim, we computed total directed interdependence between brain regions, defined as the sum of DC estimates, *S_X↔Y_* = *F_X→Y_* + *F_Y→X_* where *X* is a ROI in the left hemisphere and *Y* the homologous ROI in the right hemisphere. Figure 3c shows the distribution of the homotopic bidirectional DC, averaged over homotopic pairs of regions. DC was significantly higher in controls than LH patients, but not RH patients (*one-way Anova with group as factor, F(2,136)=3.6, p=0.03; post-hoc t-tests C vs LHP, T(84)=2.9, p=0.006; C vs RHP, T(77)=1.2, p=0.22*). Considering different networks separately (Fig. 3f), we found that for all networks, the bidirectional information flow was higher in healthy controls than LH patients (*two-way repeated-measures Anova with group and network as factors, group: F(2,1224)=3.7, p=0.03; network: F(9,1224)=75.8, p<10^-10^; interaction: F(18,1224)=1.3, p=0.14*).

We then studied whether stroke induces an asymmetry in information flow between the hemispheres by quantifying the asymmetry in information flow between brain regions, defined as the difference in DC, *G_X→Y_* = *F_X→Y_* – *F_Y→X_*. We computed the net homotopic DC asymmetry *G_X→Y_* where *X* is a ROI in the right hemisphere and *Y* the homologous ROI in the left hemisphere. *G_X→Y_* larger than zero implies a net information flow from the right to the left hemisphere, and vice versa for *G_X→Y_* smaller than zero. Figure 4a shows the distribution of the homotopic DC asymmetry, averaged over homotopic pairs of ROIs. The net homotopic information flow was shifted towards the left hemisphere in LH patients compared to control subjects, and significantly shifted towards the right hemisphere in RH patients compared to control (*one-way ANOVA with group as factor, F(2,136)=13.5, p=4·10^-6^; post-hoc t-tests C vs LHP, (T(84)=3.1, p=0.002, C vs RHP, T(77)=-2.5, p=0.02*). In other words, the DC from the intact to the lesioned hemisphere tended to be higher than in the opposite direction, implying a net information flow from the intact to the lesioned hemisphere. Considering individual RNSs (Fig. 4d), we found that for all networks, the information flow captured by the DC asymmetry was higher from the healthy to the lesioned hemisphere in patients (*two-way repeated measures ANOVA with group and network as factors, group: F(2,1224)=11.1, p=4·10^-5^; network: F(9,1224)=2.3, p=0.02; interaction: F(18,1224)=1.2, p=0.2*). Figure 4d additionally shows that the net information flow in healthy participants was preferentially from the left (dominant) towards the right (non-dominant) hemisphere, but the net asymmetry was much weaker than that observed in patients.

We then investigated whether the asymmetry effect could be attributed to a reduction of DC from the lesioned to the intact hemisphere, or rather to an enhancement of DC from the intact to the lesioned hemisphere. We analysed separately homotopic DC terms *F_X→Y_, F_Y→X_* where *X* is a ROI in the left hemisphere and *Y* the homologous ROI in the right hemisphere. The results revealed that the asymmetry is due to a reduction of DC from the lesioned to the healthy hemisphere. Indeed, we found that the homotopic DC from the lesioned to the healthy hemisphere (left to right) in LH patients was significantly lower than in healthy controls, while the DC from the healthy to the lesioned hemisphere (right to left) was only slightly, but not significantly reduced. For RH patients, the DC from the lesioned to the healthy hemisphere (right to left) was significantly lower in comparison with healthy controls, while the DC from the healthy to the lesioned was comparable (*T-test C vs LHP, left to right, T(84)=3.9, p= 3·10^-4^, right to left, T(84)=1.6, p=0.1; T-test C vs RHP, left to right, T(77)=2.2, p=0.03, right to left, T(77)=-0.05, p=0.95*). In Fig. 4e and 4f, we show the DC from left to right and from right to left for different networks, respectively. Network-level results parallelled global results.

DC from left to right was reduced in several networks for LH patients, but not RH patients, as compared to healthy controls. DC from right to left was slightly reduced in many networks for LH patients, and more severely reduced for RH patients (*three-way ANOVA with group, network and directionality - left to right vs right to left - as factors on homotopic DC, group F(2,136)=3.7, p=0.03, network F(9,2584)=74.9, p<10^-10^, directionality F(1,2584)=5.7, p=0.02, group x directionality F(2,2584)=37.8, p<10^-10^, network x directionality F(9,2584)=2.5, p=0.01*). In summary, the homotopic directed connectivity from the lesioned to the intact hemisphere in patients was reduced with respect to healthy controls. Directed connectivity from the intact to the lesioned was slightly reduced (for LH patients) or comparable with that of healthy controls. Consequently, stroke patients present an asymmetric interhemispheric information flow, going from the healthy to the lesioned hemisphere.

### Intra-hemispheric undirected functional connectivity and Granger causality analyses

We then investigated intra-hemispheric UFC, IC, and DC. Previous studies reported a bilateral increase of specific intra-hemispheric functional connections (Baldassarre et al., 2014; Eldaief et al., 2017; Ramsey et al., 2016; Siegel et al., 2016). In our analysis, we averaged over all intra-hemispheric connections and indeed observed an increase of intra-hemispheric UFC. Figure 5a shows the total (left plus right) intra-hemispheric UFC for healthy controls and stroke patients. We only observed a marginal increase for LH patients and RH patients in comparison to controls (*one-way ANOVA, F(2,136) = 2.2, p= 0.12: post-hoc T-test C vs LHP T(84)=-2.08, p=0.04, C vs RHP, T(77)=-2.24, p=0.03*). When considering different resting-state networks separately (for each network, we considered the sum of its connections with all networks), we obtained similar results (Figure 5d), with the three groups not showing strong differences (*two-way repeated-measures ANOVA with group and network as factors, group: F(2,1224)=1.5, p=0.2; network: F(9,1224)=259.3, p<10^-10^; interaction: F(18,1224)=0.8, p=0.7*).

**Figure 5:**
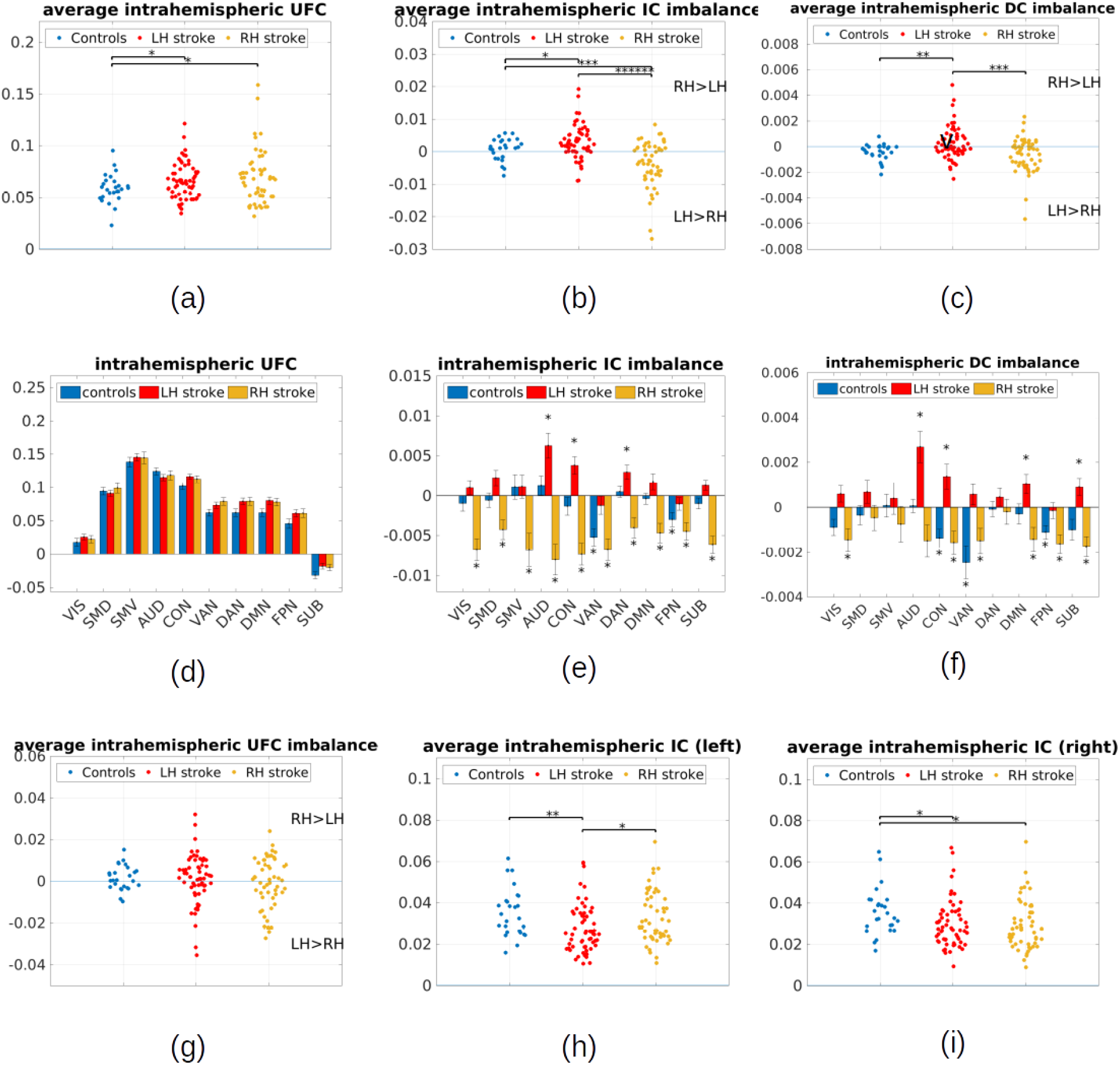
Average intra-hemispheric FC in acute phase. For each subject, we have computed the average intra-hemispheric UFC, IC and DC within the LH and RH hemisphere (a) the average intra-hemispheric UFC (sum of the LH and RH averages) is higher for patients than for controls. (b-c) the average imbalance (RH-LH difference) in intra-hemispheric IC and DC is positive for LH patients and negative for RH patients, implying that the average IC and DC are higher in the intact than the lesioned hemisphere (d) We show the average intra-hemispheric UFC by resting state network. Dots are averages over subjects, error bars standard errors over subjects. Stars represent networks for which comparison with controls (two-sample T-test) is not significant. (e-f) We show the imbalance in average intra-hemispheric IC, DC by resting state network. Dots are averages over subjects, error bars standard errors over subjects. Stars represent networks for which comparison with 0 (one-sample T-test) is not significant. (g) there is no left/right imbalance in intra-hemispheric UFC for patients and controls (h) the average intra-hemispheric IC in the left hemisphere is reduced in LH patents compared to controls and RH patients (i) the average intra-hemispheric IC in the right hemisphere is reduced in RH and LH patients compared to controls. VIS=visual, SMD=sensorimotor dorsal, SMV=sensorimotor ventral, AUD=auditory network, CON=cingulo-opercular network, VAN=ventral attention network, DAN=dorsal attention network, DMN=default mode network, FPN=fronto-parietal network SUB=subcortical nodes.

We then investigated whether stroke impacts the balance in intra-hemispheric functional connectivity between the lesioned and intact hemisphere. We computed a measure of intra-hemispheric imbalance defined as the difference between the mean UFC averaged over all pairs of regions within the same hemisphere. The intra-hemispheric UFC did not show a significant imbalance between the left and right hemisphere in either patients or controls (*one-way ANOVA, F(2,136)=0.97, p=0.38; post-hoc T-tests V vs LHP, T(84)=0.27, p=0.78, C vs RHP, T(77)=1.425, p=0.15, LHP vs RHP, T(111)=1.1, p=0.27*).

The average intra-hemispheric IC was significantly reduced in patients as compared to controls. LH patients presented a reduced intra-hemispheric IC in both hemispheres (Fig 5h and 5i), with a more pronounced reduction in the lesioned one. In RH patients, the IC was reduced only in the lesioned hemisphere. (*T-test left IC, C vs LHP: T=3.0, p=0.004, C vs RHP: T=0.7, p=0.48; T-test right IC, C vs LHP, T=2.17, p=0.03; C vs RHP T=2.31, p=0.02*). Thus, both LH and RH patients presented an imbalance in intra-hemispheric IC between the two hemispheres: intra-hemispheric IC was reduced in the lesioned hemisphere. Figure 5b shows the distribution of the intra-hemispheric IC imbalance (difference between the average intra-hemispheric IC in right and left hemisphere) for patients and controls. The imbalance is significantly different in the three groups, being significantly positive for LH patients (higher IC in the right hemisphere), significantly negative for RH patients (higher IC in the left hemisphere), and not significant for controls (*one-way ANOVA with group as factor, F(2,136)=19.6, p=3·10^-8^; post-hoc T-test, C vs LHP T(84)=-2.5, p=0.01, C vs RHP, T(77)=3.7, p=0.0004, LHP vs RHP, T(111)=5.7, p=1·10^-7^*). Figure 5e shows the intra-hemispheric IC imbalance for different resting state networks. For each network, we considered the sum of its connections with all networks. We observed a significant imbalance for both LH and RH patients in the AUD, CON and DAN networks. For RH patients, we found an imbalance also in VIS, SMD, SMV, DMN and FPN (*two-way repeated-measures ANOVA with group and network as factors, group: F(2,1224)=19.7, p<3·10^-8^; network: F(9,1224)=5.4, p=2·10^-7^; interaction: F(18,1224)=4.1, p=2·10^-8^; post-hoc T-test C vs LHP, p<0.05 FDR-corrected in AUD, CON, DAN; C vs RHP, p<0.05 FDR-corrected in AUD, CON, DAN, VIS, SMD, SMV, DMN, FPN*). The stronger effects observed in RH patients could be explained by lesion volume, since LH patients have an average wider lesions than RH patients. Healthy subjects presented a significant imbalance in the VAN and FPN (i.e., the IC was higher in the left hemisphere). In summary, stroke patients showed lower intra-hemispheric IC in the lesioned hemisphere than the intact one. Compared to healthy subjects, the intra-hemispheric IC was found to be lower in both hemispheres (more severely in the lesioned one) for LH patients and in the lesioned hemisphere for RH patients.

Finally, we analyzed DC within each hemisphere and computed a bidirectional DC strength, defined as *S_X↔Y_* = *F_X→Y_* + *F_Y→X_*. This metric was computed over all pairs of ROIs within each hemisphere. Compared to healthy controls, LH patients presented a slight reduction in intra-hemispheric DC in the left hemisphere, but not in the right hemisphere. RH patients presented no significant difference with controls (*DC left hemisphere, T-test C vs LHP, T=1.6, p=0.10; C vs RHP left: T=0.28, p= 0.77; DC right hemisphere, C vs LHP, T=0.84, p= 0.40, C vs RHP, T=0.59, p= 0.55*). All groups presented an imbalance in intra-hemispheric DC (Figure 5c). For patients, the intra-hemispheric DC was higher in the healthy hemisphere. For controls, the intra-hemispheric DC was higher in the left hemisphere, which is the dominant hemisphere. The left-ward imbalance effect appeared only slightly strengthened in RH patients compared to control subjects (*one-way ANOVA with group as factor, F(2,136)=8.12, p=5·10^-4^; post-hoc T-tests C vs LHP T(84)=-3.3, p= 0.0016, C vs RHP T(77)=1.0, p=0.34, LHP vs RHP T(111)=3.6, p=0.0005*). In Figure 5f, we show the imbalance in intra-hemispheric DC for different resting state networks. For each network, we considered the sum of its incoming and outgoing connections with all other networks. We observed a significant imbalance in CON, AUD and DMN and subcortical regions for both LH and RH patients. Additionally, RH patients had a significant imbalance in VIS, VAN and FPN networks. We observed an imbalance in VIS, VAN, FPN and the CON networks also for healthy controls, so the RH patients’ effect may not be an anomaly related to stroke (*two-way repeated-measures ANOVA with group and network as factors, group: F(2,1224)=8.9, p=0.0002; network: F(9,1360)=3.7, p=0.0001; interaction: F(18,1224)=2.6, p=0.0002; T-test C vs 0, p < 0.05 FDR-corrected for VIS, VAN, FPN, CON; LHP vs 0, p < 0.05 FDR-corrected for CON, AUD, DMN; p<0.05 FDR-corrected for VIS, VAN, FPN*).

### Global FC and GC summary measures and stroke-related behavioral deficits

In the previous sections, we characterized several global correlates, or summary measures, of stroke based on functional connectivity and Granger causality analyses. Four summary measures were related to homotopic connections: 1) UFC_homo_: homotopic UFC; 2) IC_homo_: homotopic IC; 3) ΣDC_homo_: sum of homotopic DC (contralesional to ipsilesional plus ipsilesional to contralesional); 4) ΔDC_homo_: homotopic DC asymmetry (contralesional to ipsilesional minus ipsilesional to contralesional). Both LH and RH patients, as a group, presented a reduced UFC_homo_, a reduced IC_homo_ and an enhanced ΔDC_homo_ in comparison to healthy subjects. In addition, LH patients presented a reduction of ΣDC_homo_. Three additional summary measures were related to intra-hemispheric connections: 5) ΣUFC_intra_: sum of intra-hemispheric UFC (ipsilesional plus contralesional); 6) ΔIC_intra_: intra-hemispheric IC imbalance (contralesional minus ipsilesional); 7) ΔDC_intra_: intra-hemispheric DC imbalance (difference of contralesional minus ipsilesional). LH and RH patients present an enhanced ΣUFC_intra_ and an enhanced ΔIC_intra_ in comparison to healthy subjects.

In order to study whether these global summary measures were correlated among each other, we computed the partial Spearman correlation values between pairs of measures for all patients, controlling for lesion volume. Results are shown in Fig. 6a. We observed that the measures split into three groups. The first group included UFC_homo_, IC_homo_ and ΣDC_homo_, which were all strongly correlated. These three measures quantified the strength of inter-hemispheric (homotopic) connectivity. The second group included ΔDC_homo_, ΔIC_intra_ and ΔDC_intra_, which were mutually correlated and uncorrelated with the homotopic measures. These three measures quantified homotopic imbalance. Last, ΣUFC_intra_ was weakly correlated with the other measures. A PCA on the (z-scored) summary measures identified two principal components explaining 32% and 30% of the total variance respectively (Fig. 6b), henceforth indicated as the principal components PC1 and PC2. The PC1 loaded on UFC_homo_, IC_homo_ and ΣDC_homo_, whereas the PC2 on ΔDC_homo_, ΔIC_intra_ and ΔDC_intra_. Intuitively, PC1 summarized the inter-hemispheric functional integration (Fig. 6b, left panel), whereas PC2 the inter-hemispheric imbalance (Fig. 6b, right panel). We investigated whether PC1 and PC2 were related to the structural lesions. As shown in Fig. 6c, PC1 was negatively correlated with lesion volume (*Spearman r=-0.47, p<10^-6^*): the larger the lesion, the lower the functional integration between the hemispheres (Fig. 6c). Concerning PC2, we found that the modulus of PC2 was positively correlated with lesion volume (*Spearman r=0.54, p<10^-7^*): the larger the lesion, the larger the asymmetry between the hemispheres (Fig. 6d). In this regard, we should note that the value PC2 reflected the direction of the asymmetry (left-ward or right-ward), while its modulus reflected the magnitude of the asymmetry.

**Figure 6.**
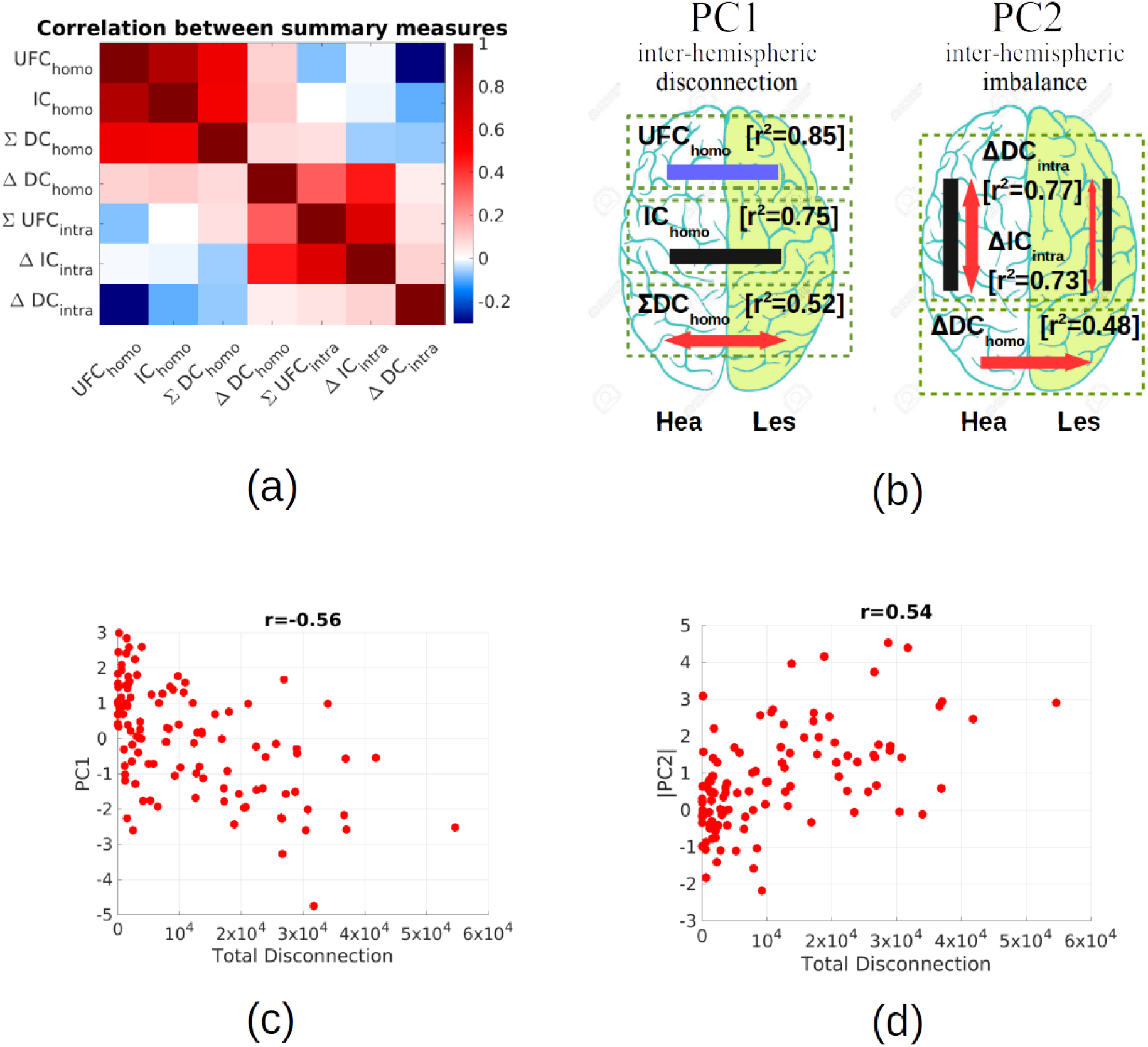
Global FC and GC summary measures. Our analysis identified several global correlates of stroke, or summary measures, based on functional connectivity and Granger causality analyses. (a) Looking at the correlation (partial Spearman correlation correcting for lesion volume) between each pair of measures, one can immediately notice two separate groups of correlated measures, one including UFC_homo_, IC_homo_, ΣDC_homo_, the other including ΔDC_homo_, ΔIC_intra_, ΔDC_intra_. (b) A PCA on the seven measures revealed two PCs explaining more than 32% and 30% of the total variance across patients. The first component (PC1) loaded on UFC_homo_, IC_homo_, ΣDC_homo_, the second component (PC2) on ΔDC_homo_, ΔIC_intra_, ΔDC_intra_. This is summarized in the two brain plots showing intuitively the main effects captured by PC1 and PC2 in the healthy and lesioned hemisphere. (c) PC1 correlates negatively with lesion volume (d) the modulus of PC2 correlates positively with lesion volume.

In a previous work (Corbetta et al., 2015), eight behavioral scores were identified, corresponding to the eight strongest principal components explaining a large fraction of variance in behavioral tests covering language, memory, motion and attention function. The eight factors were associated with language, left body motion, right body motion, spatial attention (hemispatial neglect), sustained attention, shifting attention, spatial memory, verbal memory. Higher scores signify better performance. Right body motion, language, verbal memory and shifting attention scores tend to be lower for LH patients, sustained attention scores show no hemispheric bias, while left body motion and spatial memory scores tend to be lower for RH patients.

We then quantified to which extent the first two principal components (PC1 and PC2) were predictive of the observed behavioral deficits. To do so, we computed the Spearman correlation between the two PCs and behavioral scores (Fig. 7a). We had two general predictions. First, we expected a positive correlation between performance and inter-hemispheric integration (Carter 2010, Siegel 2016, Corbetta 2018). Consequently, we expected PC1 to correlate positively with behavioral scores. Second, we expected that a decrease of connectivity within and from the lesioned hemisphere would have been generally detrimental for performance. Hence, behavioral scores were expected to correlate negatively with PC2. These expectations were partially met. PC1 correlated positively with all scores for both LH and RH patients (*LHP: Spearman r> 0.17 for all scores, RHP: Spearman r > 0.2 for all scores except Shift Att*). It is noteworthy that different correlations were significant in LH and RH patients. For LH patients, effects were stronger for scores related to verbal function, general attention, and contralesional motion (*r>0.35, p<0.05 for Lang, Ver Mem, Shift Att, Sust Att, Mot L, FDR-corrected for 8 comparisons)*. For RH patients, effects were stronger for scores related to spatial processing and motion (*r> 0.47, p<0.05 Sp Mem, Sp Att, Mot L, Mot R, FDR-corrected for 8 comparisons*). PC2 correlated negatively for LH patients (*Spearman r < −0.15, p<0.31 uncorrected for all scores except Shift Att),* and was significant for scores associated with language (*p<0.05 Lang, Ver Mem, FDR-corrected for 8 comparisons*), while correlations were weaker, ambiguous in sign, and not significant for RH patients (*p>0.05 for all scores, FDR-corrected for 8 comparisons);* the strongest observed effect was a positive correlation between PC2 and language-related scores (*r=0.15, p=0.29 for Lang; r=0.35, p=0.03 uncorrected for Ver mem*). In summary, a lower PC1 (lower inter-hemispheric integration) was associated with significantly worse performance in several behavioral domains, different for LH and RH patients; for RH patients, the effect could not be explained by lesion volume. A higher PC2 (higher inter-hemispheric imbalance) was associated with worse performance in language-related domains for LH patients, and with better performance in RH patients.

**Figure 7.**
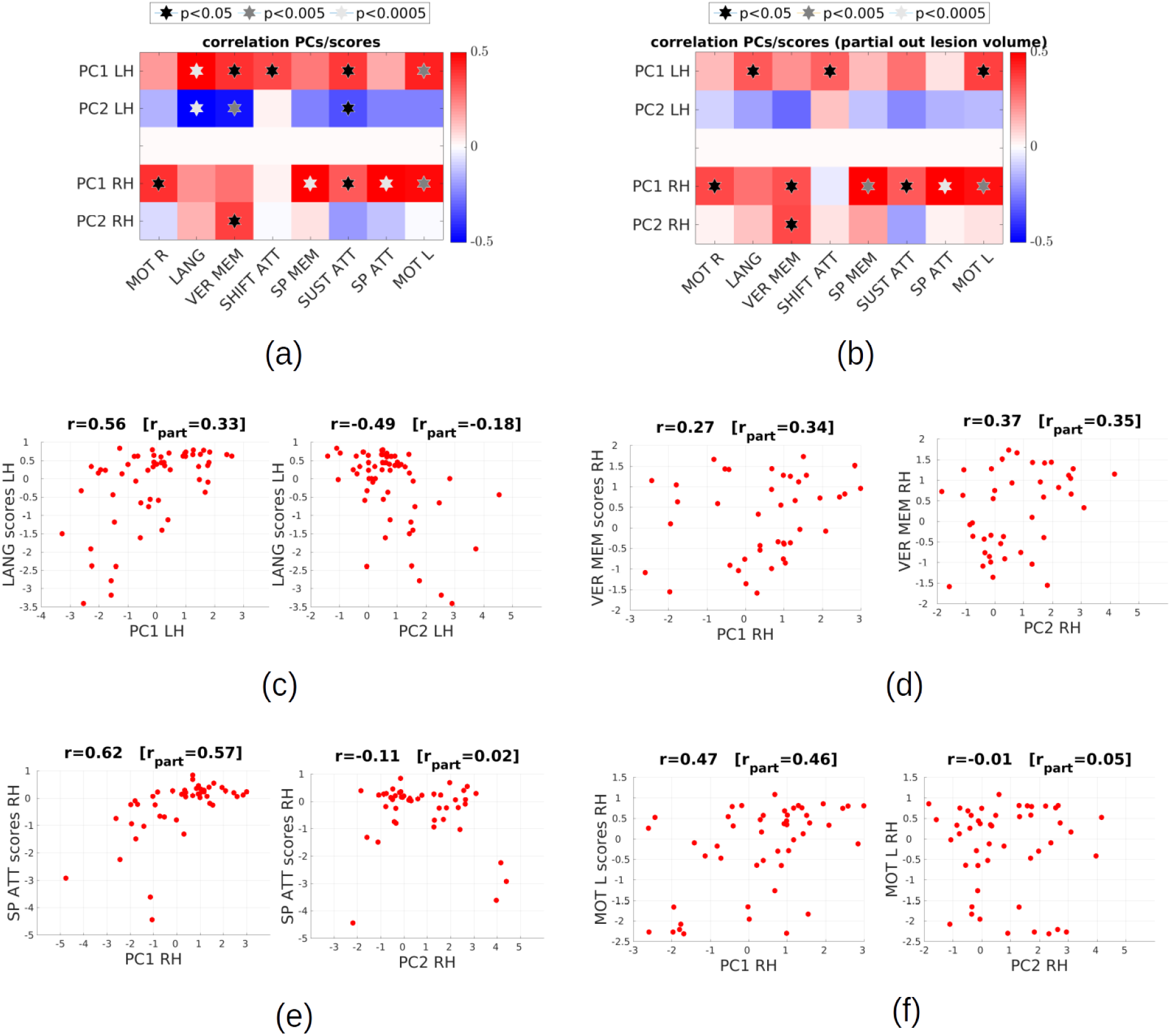
Correlation with behavioral scores.: (a) Spearman correlation between behavioral scores and the two principal components (PC) summarizing FC and GC stroke summary measures (b) partial Spearman correlation between behavioral scores and the two principal components, correcting for lesion volume (c) scatter plot of PC1/PC2 versus language scores for LH patients (d) scatter plot of PC1/PC2 vs verbal memory scores for RH patients (e) scatter plot of PC1/PC2 vs spatial attention scores for RH patients (f) scatter plot of PC1/PC2 versus left body motion scores for RH patients

### Control analyses

Part of the observed differences between LH and RH patients may be due to the influence of lesion volume. Both the homotopic UFC and the homotopic IC appeared to be higher for RH than LH patients (Fig. 3). In both cases the difference could be largely explained by differences in lesion volume (*UFC: one-way ANOVA with group as factor, after linearly regressing logarithm of lesion volume lesion: F(1,1110) = 1.5, p= 0.2; IC: one-way ANOVA with group as factor, after linearly regressing logarithm of lesion volume lesion: F(1,1110)=3.4, p=0.06*). Moreover, since PC1 and PC2 correlate with lesion volume, part of the observed correlation with behavioral scores may be explained by lesion volume. A larger lesion volume is causally related to more widespread structural disconnections (Griffiths et al. 2019), which are at the root of functional connectivity alterations captured by PC1, PC2. However, a larger lesion volume is also causally associated with a larger local damage to cortical area. Thus, a correlation between the PCs and behavioral scores does not provide sufficient evidence that the functional anomalies captured by the PCs have a specific role in the genesis of the deficits, more than other functional perturbations such as impaired activity in the locally damaged area. We computed the partial Spearman correlation between the two PCs and behavioral scores, controlling for the effect of lesion volume (Fig. 7b). As for PC1, we still observed a positive correlation with behavioral scores (*LHP: Spearman r> 0.13 for all scores except Sp Att; RHP: r > 0.26, for all scores except Shift Att*). Surprisingly, while effects were generally reduced and no longer significant for LH patients (*p>0.05 for all scores, FDR-corrected for 8 comparisons),* correlations remained significant for RH patients (*r> 0.36, p<0.05 Mot R, Sp Mem, Sp Att, Mot L, FDR-corrected for 8 comparisons*). For PC2, we still observed a generally negative correlation for LH patients (*Spearman r < −0.08, for all scores except Shift Att),* but correlations were no longer significant (*p>0.05 for all scores, FDR-corrected for 8 comparisons*). Correlations were still not significant for RH patients (*p>0.05 for all scores, FDR-corrected for 10 comparisons*).

We finally performed control analyses to investigate potential confounding effects associated with nuisance sources and hemodynamic lags. GC analyses were performed on preprocessed BOLD signals without global signal regression (GSR) removal. The rationale for this choice was that GSR may effectively work as a “temporal filter” (Liu et al., 2017, Nalci et al., 2019), suppressing the contribution of time points associated with low global signal, potentially distorting the estimation of information flows in GC. While standardly adopted for UFC estimation, GSR is a contentious step (Saad et al., 2012), particularly when one compares healthy subjects with neurological or psychiatric patients (Hahamy et al., 2014; Yang et al., 2014). Indeed, the global signal can reflect extended correlation of neural origin (Schölvinck et al., 2010), possibly differing between patients and control subjects. By applying GSR our data, homotopic information transfer (homotopic IC and bidirectional DC) presented similar effects to those found without GSR, including the asymmetry in homotopic DC (Fig. 8a). However, results on intra-hemispheric GC differed: no clear imbalance is observed in intra-hemispheric DC or IC (Fig. 8b and 8c). Thus, GSR significantly attenuates the hemispheric imbalances. However, due to the high network specificity of the observed imbalances, it appears unlikely that such imbalances represent metabolic, movement, breathing-rate, cardiovascular or vigilance effects. It is more likely that differences in global signal between the hemispheres represent alterations in the excitation/inhibition balance within each hemisphere (Yang et al., 2014), which are obscured by GSR.

**Figure 8.**
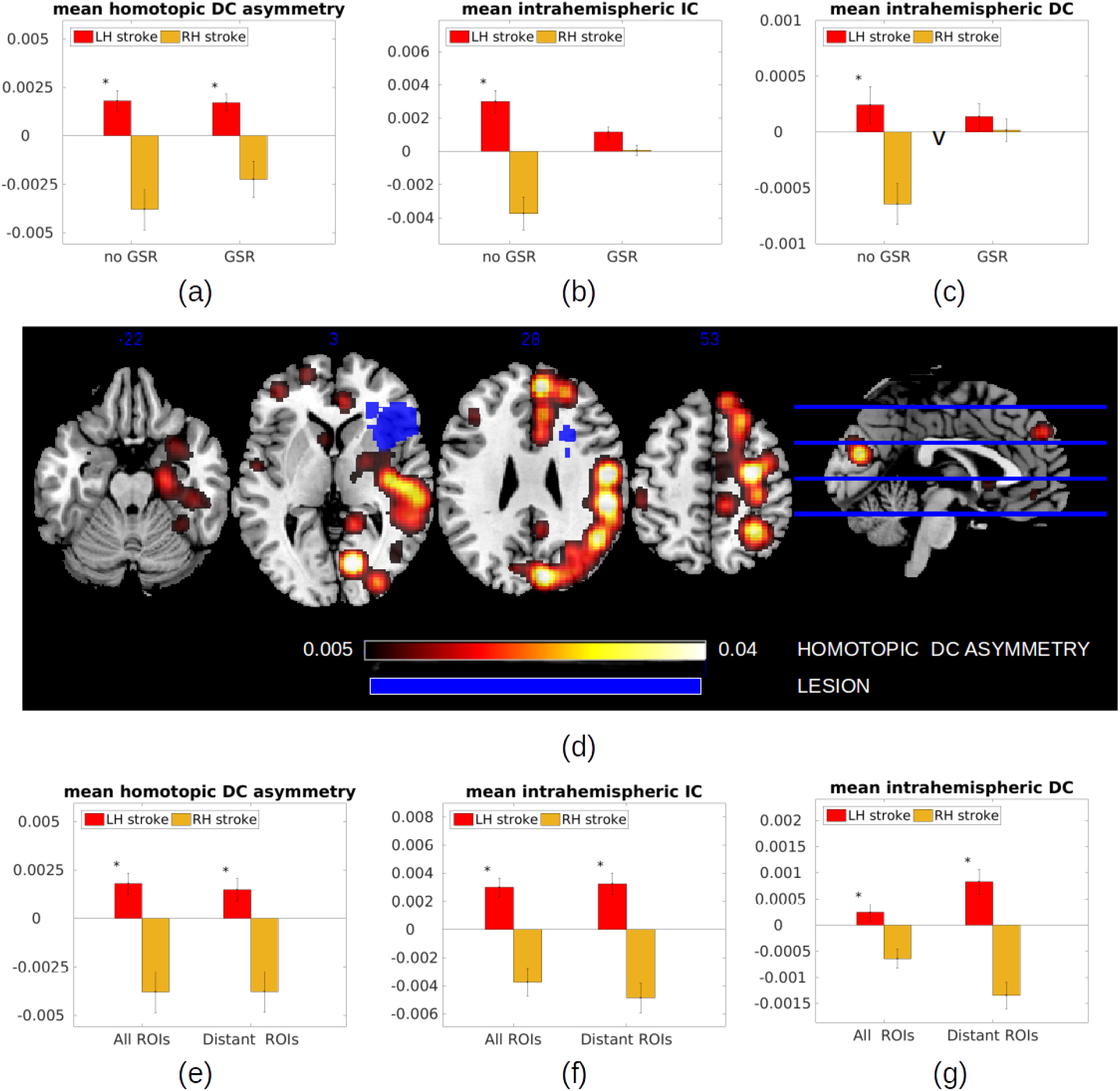
Control for possible confounds. (a-c) We checked the effect of GSR on the found inter-hemispheric imbalances. GSR has no effect on the homotopic DC asymmetry, while it removes the imbalance in intra-hemispheric IC and DC. (d-g) We checked possible influences of perilesional hemodynamic anomalies on our results. In (d) we verified whether the homotopic DC asymmetry could be driven by hemodynamic lags in the perilesional area. We show a map of the homotopic DC asymmetry for one representative subject, together with the lesion location (in blue). Strongest homotopic DC asymmetries are found far from the lesion. In (e-g) we show the effect of removing from analysis all regions at a distance < 4cm from the lesion. Such removal has no effect on the homotopic DC asymmetry, while it strengthens the imbalance observed in intra-hemispheric IC and DC.

Hemodynamic lags represent an additional potential confound for our results. In fact, stroke can cause a pathologic delay in the hemodynamic response in the perilesional area, or in a wider area subserved by the occluded artery (Siegel et al., 2016b). This delay may introduce spurious “lags” of non-neural origin between regions in this area and homologous regions in the intact hemisphere, thus contributing to the observed homotopic DC asymmetry. We checked whether the observed global homotopic DC asymmetry could be linked to asymmetries in the perilesional area. We considered each region *X* in the lesioned hemisphere and computed the DC asymmetry *G_Y→X_* = *F_Y→X_* – *F_X→Y_* where *Y* is the homologous area in the intact hemisphere. We thus obtained brain-wide maps of homotopic DC asymmetry that overlayed with the lesion maps (to produce the homotopic DC maps, we assigned the value *G_Y→X_* to all voxels within a radius of 10mm around the center of each ROI *X,* and then applied 10mm Gaussian smoothing). In Fig. 8d, we show the results for a representative subject.The strongest DC asymmetries were observed far from the lesions location in the brain. In order to have a more quantitative control, we repeated our analyses excluding all regions at a distance less than 4cm from the lesioned area. As shown in Fig. 8e-8g, the homotopic DC asymmetry is still present after this removal, while the intra-hemispheric IC and DC imbalance appear to be even strengthened. This showed that the observed effects are not due to anomalous hemodynamic lags in the vicinity of the lesion.

## DISCUSSION

Previous research, including previous work on the Washington stroke database (Joshua Sarfaty Siegel et al., 2016, Corbetta et al., 2018, Griffis et al. 2019), has made extensive use of resting-state fMRI to investigate functional connectivity in stroke patients. The results of these studies suggest a view of stroke as a network dysfunction syndrome. Stroke is accompanied by widespread alterations of functional connectivity, with common patterns observed across patients independently of lesion location. In particular, most patients present a loss of inter-hemispheric FC (Corbetta et al., 2018; Golestani et al., 2013; Joshua Sarfaty Siegel et al., 2016; Tang et al., 2016). Anomalies of long-range FC are paralleled by perturbations of monosynaptic (Griffis et al. 2019) and polysynaptic (Griffis et al. 2020) structural connections. While sensorimotor deficits are reasonably well explained by local damage, cognitive deficits are better explained by network dysfunction (Siegel et al., 2016).

However, it was still unclear whether stroke produces functional asymmetries in long-range brain interactions. Being symmetric, standard functional connectivity (UFC) and structural connectivity (SC) measures are not suited to address this question. In our work, we used Granger causality (GC) measures to investigate alterations of long-range directional interactions in the brain after stroke. By exhaustively looking at temporal dependencies between the BOLD signals of two regions, GC measures can yield information about the directionality and time scale of interactions, which is missing from UFC analyses. Our analyses revealed several stroke-related asymmetries between the hemispheres, which further allowed us to better highlight major differences between patients with left- or right-hemisphere lesions which had not been specifically addressed in previous analyses.

### Stroke-related modulations in inter- and intra-hemispheric coupling revealed by Granger causality analyses

One of the major functional effects of stroke is a loss of inter-hemispheric integration associated with a decrease of homotopic UFC. It is still relatively unknown to which extent the UFC decrease corresponds to a decrease of direct interactions (supported by homotopic connections crossing the corpus callosum (Schmahmann et al., 2009)) or indirect interactions through subcortical structures. Our results from GC-based analyses show that UFC decrease (Fig. 3a) is strongly associated with a loss of inter-hemispheric interactions captured by the homotopic IC and DC (Fig. 3b, 3c). IC captured cortico-cortical interactions unfolding within 1 TR, while DC captured lagged cortico-cortical interactions occurring on a time scale longer than 1 TR. Classically, IC are interpreted as originating from external common inputs (Ding et al., 2006). Since a large part of the total interdependence between the signals of homotopic areas is due to the IC (Fig. 2c), our results suggest that a component of stroke-related alterations in cortico-cortical coupling emerges from disrupted common inputs from regions that project symmetrically to cortical areas, such as subcortical structures. This hypothesis is supported by structural analyses that locate stroke lesions primarily in subcortical areas, such as the thalamus (Corbetta et al. 2015), as well as by recent experimental work showing that subcortical structures can play a large role in maintaining FC between cortical regions when direct influences are impaired (Canella et al. 2020). However, given the slow sampling rate of our data (TR=2s), an IC decrease cannot be uniquely attributed to a loss of common input, as it may also result from a decrease of fast directed interactions occurring on timescale shorter than 2s. To which extent subcortical structures contribute in re-modulating cortical interaction remains a relevant topic for further investigation.

Importantly, even though IC rather than DC dominates homotopic interdependence, DC analysis is precious as it hints at strong interhemispheric communication asymmetries. Our results on homotopic DC showed that stroke impacts the inter-hemispheric information flow asymmetrically, with a spared information flow from the healthy to the lesioned hemisphere and a reduced flow in the opposite direction (Fig. 4a). This asymmetric effect is not immediately explained by structural lesions, since there is no evidence that ischemia would affect selectively fibers from the ipsilesional to the contralesional hemisphere rather than in the opposite direction. The homotopic asymmetry we measured is in line with recent work (Wang 2019) showing that time series in the lesioned hemisphere are “lagged” with respect to the homologous areas in the healthy hemisphere. To which extent this effect may stem from non-neural, hemodynamic causes - a systematic alteration of the hemodynamic response in the lesioned hemisphere - remains an open question. In our control analysis (fig. 8) we excluded the possibility that the effect be trivially related to the well-known presence of large hemodynamic lags in the perilesional area (Siegel et al., 2016b). Thus, the measured asymmetry, if caused by hemodynamic effects, would imply wide alterations of the hemodynamic response far from the lesion. In future work, this issue may be specifically addressed by applying deconvolution prior to GC analysis, building on “blind deconvolution” techniques that allow retrieving the hemodynamic response from resting-state data (Wu et al., 2013). Such analysis is beyond the scope of the current study. Since deconvolution techniques for resting-state fMRI remain exploratory, it is still unclear whether these methods are accurate in presence of anomalous distortions of the hemodynamic response potentially arising in pathological conditions such as stroke. The relevance of the observed homotopic GC asymmetry is strengthened by our analysis of intra-hemispheric GC, which revealed another functional imbalance between the hemispheres in stroke patients: intra-hemispheric IC and DC are higher in the intact hemisphere than the lesioned one (Fig. 5b and 5c). Our results are not conclusive regarding the relation between the homotopic DC asymmetry (Fig. 4a) and the imbalance in intra-hemispheric IC and DC (Fig. 5). However, we provided evidence that the intra-hemispheric and inter-hemispheric imbalances are correlated (Fig. 6), which suggests that the two results are not independent and may have a common cause. We speculate that both effects could stem from structural disconnection within the lesioned hemisphere, causing a loss of inter-areal excitatory influences. Since stroke can damage structural connections between ipsilesional areas, we could generally expect a loss of excitatory influences, and hence general activity decrease, within the lesioned hemisphere (Grefkes and Fink, 2014). This, in turn, would also imply that the lesioned hemisphere would exert less excitation on the healthy one. This picture would explain both the decrease of ipsilesional DC and IC, and the decrease of DC from the lesioned to the healthy hemisphere. Further support to this interpretation comes from the fact that all imbalance measures (ΔDC_homo_, ΔIC_intra_, ΔDC_intra_) correlate negatively with lesion volume (i.e., the stronger the lesion, the higher the intra- and inter-hemispheric functional imbalances).

Post-stroke inter-hemispheric imbalances in effective connectivity were widely reported in the motor system, as reviewed in (Grefkes and Fink, 2014). During motor tasks, excitatory influences within the lesioned hemisphere are reduced, contributing to a general decrease of ipsilesional brain activity (Grefkes and Fink, 2014; Rehme and Grefkes, 2013). As for inter-hemispheric connectivity, several studies on the motor system after stroke indicate an anomalous influence of the contralesional hemisphere onto the lesioned one during motor tasks (Rehme and Grefkes, 2013, Grefkes et al., 2010, Grefkes and Fink, 2014). Whether the contralesional influence is inhibitory (hence detrimental to motor performance), or excitatory (hence supportive of performance) seems to depend on several factors, including time after stroke and severity of the lesions (Pino et al., 2014). Our results instead showed a decrease of influence of the damaged hemisphere on the normal one. However, we are wary of a direct comparison, since our whole-brain results were obtained with a resting-state paradigm, hence without any specific involvement of the motor cortex. In order to further clarify inter-hemispheric balance after stroke, future whole-brain studies should discriminate between excitatory and inhibitory influences, which is not possible in the current GC analysis.

### Hemispheric functional imbalance and stroke-related behavioral deficits

Previous behavioral analyses on this cohort (Corbetta et al., 2015; Ramsey et al., 2017) identified sets of correlated deficits for left and right lesions respectively, largely agreeing with lateralization maps described in healthy subjects (Karolis et al., 2019). Right body motion, language, verbal memory and shifting attention scores tend to be lower for LH patients, sustained attention scores show no hemispheric bias, while left body motion, spatial attention and spatial memory scores tend to be lower for RH patients. Here, we systematically addressed for the first time the functional basis of LH/RH patient differences.

In LH patients, the amount of inter-hemispheric communication (summarized by PC1) correlated positively with behavior for domains that are specific to the left hemisphere (language, verbal memory, attention shifting), with the exception of right motion. The correlation between PC1 and behavioral scores was significantly lower if lesion volume was regressed, and presently we cannot discriminate the specific impact of inter-hemispheric communication loss on behavioral function from other possible effects resulting from the lesion. The imbalance between the hemispheres (summarized by PC2) correlated negatively with behavioral scores, significantly for behaviors that were affected by left hemisphere lesions (language, verbal memory) and could be largely explained by lesion volume, suggesting that it reflects the extent of intra-hemispheric LH damage.

In RH patients PC1 correlated with behavior for domains more associated with the right hemisphere (motor function, spatial and sustained attention, and spatial memory). Correlations were robust to regression of lesion volume, which suggests a specific impact of inter-hemispheric communication loss on behavior. This hypothesis agrees with previous studies showing that deficits that were affected by right lesions were more associated with inter-hemispheric rather than intra-hemispheric functional disconnection (Baldassarre et al., 2016a, 2016b; Siegel et al., 2016). We speculate that input from the LH may be more critical for functional integrity of the RH than the other way around, congruently with studies reporting that the left hemisphere presents more central or indispensable regions for the whole-brain structural network (Iturria-Medina et al., 2011), and that the right hemisphere depends more heavily on integration with the left one than the other way around (Gotts et al., 2013). In RH patients we did not observe a negative correlation between scores and PC2, suggesting that intra-hemispheric damage has a lesser impact on behavior. Instead, we observed a positive correlation between PC2 and verbal memory scores, which suggests a supportive role of the left (contralesional) hemisphere for a left-lateralized function in the case of right lesions.

In both LH and RH patients, sustained attention scores had a significant positive correlation with PC1, and a negative correlation with PC2. This is consistent with previous literature suggesting that higher scores are associated with a higher inter-hemispheric integration and a higher intra-hemispheric integration in the lesioned hemisphere (Corbetta et al. 2005, He et al. 2007, Corbetta and Shulman 2011), but a large part of the correlation observed in this work could be explained by lesion volume, hence at present we cannot know to which extent the effect is causally related to functional connectivity anomalies.

### Methodological considerations on Granger causality analyses: advantages and limitations

The efficacy of GC as a data-driven analysis method rests on its ability to uncover global patterns of information flow and differences in information flows between groups or experimental conditions in a completely unsupervised way (Faes et al., 2017; Friston et al., 2013; Roebroeck et al., 2011, 2005). The pairwise covariance-based GC analysis approach used in this work is a method of choice for whole-brain analyses of large databases aimed at uncovering general information flow patterns (see e.g., Deco at al., 2021). The main limitation of our analysis is the uncertainty affecting GC estimates at different levels, from single-session to single-subject (Fig. 2). For each GC-based stroke summary measure (e.g., the total homotopic IC), we obtained a large group variance, and consequently a large overlap between the distributions of patients and controls, so that we could not robustly classify an individual as patient or control based on his/her value of the summary measure. It is likely that part of this variance reflects estimation error, rather than true interindividual variability. Analogously, the uncertainty affecting single-subject estimates also implies a difficulty in relating individual GC results with individual behavioral scores. Thus, estimation error limits the use of GC for the development of personalized biomarkers predictive of clinical condition and behavioral performance at the single-patient level. This limitation is not inherent in GC per se, but depends on the relative paucity of functional data available for each patient, and the poor temporal resolution implied by TR=2s. By taking longer recordings or repeating recording sessions, we could obtain much more accurate GC estimates. Improved GC estimates may also be obtained by using a lower TR. Using a TR=0.67s (as in the Human Connectome Project database (van Essen et al., 2013)) would triple the number of points for estimation and offer a significantly improved time resolution, allowing for a more precise characterization of directionality effects (we predict that by using a shorter TR, a part of the total interdependence that is seen as IC in this study would appear as DC). In our opinion, the main limitation of GC in this study is due to intrinsic properties of the data, rather than the specific approach used for calculating DC - bivariate and covariance-based. We are skeptical that more sophisticated approaches for GC estimation would yield radically improved results. In particular, for several reasons we do not believe that a multivariate approach (Barnett 2014), which in principle gives cleaner results by eliminating indirect network effects, would particularly contribute to our study. Since we have a large number of areas, conditioning would impair estimation, especially because of many redundancies (Stramaglia et al. 2016). Moreover, since our main results are based on large averages over many pairs of regions, they are largely indifferent to whether single links are affected by indirect contributions.

### Conclusions

To conclude, the Granger causality (GC) analysis of inter-areal interactions after stroke highlighted two broad pathological features. First, a decrease of homotopic GC, suggesting a large decrease of interhemispheric communication, either direct or mediated by subcortical structures. Second, an inter-hemispheric imbalance, revealed by an asymmetry in homotopic GC, as well as a right-left difference in intra-hemispheric GC, suggesting a decrease of communication within and from the lesioned hemisphere. These results show that previously observed FC alterations in stroke are related to broad changes in inter-areal communication. Furthermore, our analysis confirms and generalizes previous findings about post-stroke inter-hemispheric imbalances in the motor and attention system. The observed GC anomalies highlighted a different impact of lesion on behavior depending on which hemisphere was lesioned. Left-lateralized behavior was strongly affected by loss of intra-hemispheric communication in patients with left hemisphere lesions. Right-lateralized behavior was strongly affected by loss of inter-hemispheric communication in patients with right hemisphere lesions.

## Abbreviations

fMRI: functional magnetic resonance imaging
FC: functional connectivity
RSNs: resting-state networks
GC: Granger causality
IC: instantaneous (Granger) causality
DC: directed (Granger) causality
UFC: undirected functional connectivity
LH/RH: left/right hemisphere

## Acknowledgements

MA, AB, CF and MC were supported by FLAG ERA II “Joint Transnational Call 2017” - HBP - Basic and Applied Research 2, Brainsynch-Hit (ANR-17-HBPR-0001) during completion of this work. The Authors thank Gustavo Deco and Reiner Goebel for helpful comments and suggestions during completion of this work.

## Data and code availability statement

The full set of neuroimaging data (along with behavioral data) are available at http://cnda.wustl.edu/app/template/Login. The Granger Causality code used in this work is available at https://github.com/micheleallegra/CovGC

